# EthoPy: Reproducible Behavioral Neuroscience Made Simple

**DOI:** 10.1101/2025.09.08.673974

**Authors:** Alexandros Evangelou, Maria Diamantaki, Konstantina Georgelou, Zoi Drakaki, Lydia Ntanavara, Gerasimos Gerardos, Sofia Morou, Nikolaos Chatziris, Zoi Dogani, Elissavet Anna Petsalaki, Odysseas Nikolaos Raos, Anastasios Gratsakis, Athanasia Papoutsi, Emmanouil Froudarakis

**Author notes:** Equal contributions (Alexandros Evangelou, Maria Diamantaki).

## Abstract

As brain activity is tightly coupled to behavior, an accurate understanding of neural function necessitates consideration of behavioral tasks that capture the complexity and variety animals encounter. Nevertheless, animal training for behavioral experiments is often labor-intensive, costly, and difficult to standardize. To overcome these challenges, we developed EthoPy, an open-source, Python-based behavioral control framework that integrates stimulus presentation, hardware management, and data logging. EthoPy supports diverse behavioral paradigms, stimulus modalities, and experimental systems, from homecage to head-fixed configurations, while operating on affordable hardware, such as Raspberry Pi. Its modular architecture and database integration enable scalable, high-throughput automatic behavioral training with minimal experimenter involvement while ensuring reproducibility through comprehensive metadata tracking. By automating training workflows, EthoPy makes it feasible to implement sophisticated behavioral paradigms that are traditionally difficult to achieve. EthoPy thus provides an accessible, extensible framework to study behavior and the underlying neural activity.

## Introduction

Understanding how neural activity drives behavior has advanced significantly through recent high-yield recording techniques (Jun et al., 2017; Steinmetz et al., 2021; Lakunina et al., 2025; Sofroniew et al., 2016; Zhang et al., 2024; Guo et al., 2023) and the development of ethologically relevant behavioral paradigms in animal models (Krakauer et al., 2017; Peters et al., 2015; Vielle, 2025; Shemesh and Chen, 2023). However, a key challenge remains: establishing automated and scalable behavioral tasks across diverse experimental paradigms, from homecage to head-fixed settings, and from simple port selection to complex navigation tasks (Nasr et al., 2022; Castelhano-Carlos et al., 2017; Watson et al., 2019; Nourizonoz et al., 2020; Kim et al., 2019; Lopes et al., 2015). Many behavioral experiments have limited scalability as they require intensive manual effort for animal handling and response tracking during training (Sukoff Rizzo and Silverman, 2016; Zoccolan and Di Filippo, 2018). Such involvement can also introduce variability, increase animal stress (Chesler et al., 2002; Sorge et al., 2014; Balcombe et al., 2004) and reduce reproducibility across labs (Richter, 2020; Nigri et al., 2022). Furthermore, observer-based assessments are time-consuming and prone to error, limiting the efficiency and consistency of behavioral research (Gulinello et al., 2019).

A second challenge is the lack of flexible systems that can seamlessly integrate both commercial and custom-built hardware. Existing platforms are often rigid and task-specific, restricting their adaptability to new experimental demands or emerging technologies. Ideally, behavioral systems should decouple software from hardware dependencies, allowing researchers to prototype and deploy new tasks without initiating new projects from scratch. Third, behavioral frameworks typically assume fixed task parameters throughout a session, overlooking dynamic changes in behavioral state. Yet, performance can fluctuate due to learning, motivation, or arousal (Berditchevskaia et al., 2016; Hulsey et al., 2024; Pisupati et al., 2021). This highlights the need for adaptive in real-time task designs that adjust parameters, such as difficulty, based on behavior. This design also permits closed loop approaches that enable causal control through the online task modulation (Watson et al., 2019; Kim et al., 2019; Lopes et al., 2015; Schweihoff et al., 2021). While fine control of task structure is available in many systems (Akam et al., 2022; Saunders et al., 2019), making such adaptive strategies feasible and scalable remains a key challenge.

Finally, the growing complexity of behavioral experiments has increased the need for efficient data and metadata management. Reproducibility requires careful logging of metadata, yet current practices often vary across platforms, limiting protocols and data reuse and sharing. Inconsistent access to datasets across teams further ham-pers collaboration and transparency. The FAIR data principles (Findable, Accessible, Interoperable, and Reusable (Martone, 2024)) offer a valuable framework to address these challenges. Applying the FAIR principles to behavioral neuroscience ensures that data are securely stored, clearly documented, and easily reused, supporting open science and improving long-term accessibility across diverse research groups.

Recent years have seen progress toward addressing these challenges through software tools that interface with diverse hardware components. Automated behavioral chambers now enable animals to interact with levers/buttons, sensors, or touchscreens, working towards reducing variability and improving experimental control (Watson et al., 2019; Akam et al., 2022; Saunders et al., 2019; Ameen-Ali et al., 2012; Poddar et al., 2013; Aoki et al., 2017; Gordon-Fennell et al., 2023; Iman et al., 2021). However, many systems still require manual handling and are tied to specific tasks, limiting extensibility (Watson et al., 2019; Nourizonoz et al., 2020; Kim et al., 2019; Aoki et al., 2017). Most lack rich stimuli manipulation or integration with data management tools, while only a few may support secure, scalable databases (Poddar et al., 2013). Overall, these tools still do not provide a unified, opensource solution to the identified bottlenecks, and their sustainability and uptake critically depend on their accessibility and integration to the active research community. Current behavioral research can greatly benefit from easily shareable and reproducible behavioral paradigms, remote and automatic control, supported by open access code.

## Results

### A unified control framework for behavioral experiments

We developed EthoPy, an open-source Python framework for high-throughput behavioral experiments, so it can manage all aspects of an experiment, such as design, behavior, hardware control, stimulus presentation, and data management, while prioritizing flexibility to adapt to emerging technologies and paradigms. Its modular architecture separates experimental components into independent modules (Fig. 1a), enabling easy switching between setups (homecage, head-fixed, open arena), stimulus modalities (olfactory, visual, auditory), and paradigms (Go/no-Go, 2 alternative forced choice) without rewriting code. Extensibility is achieved through abstracted core functions, allowing rapid customization and integration of new modules or community tools (Fig. 1b).

**Fig. 1.**
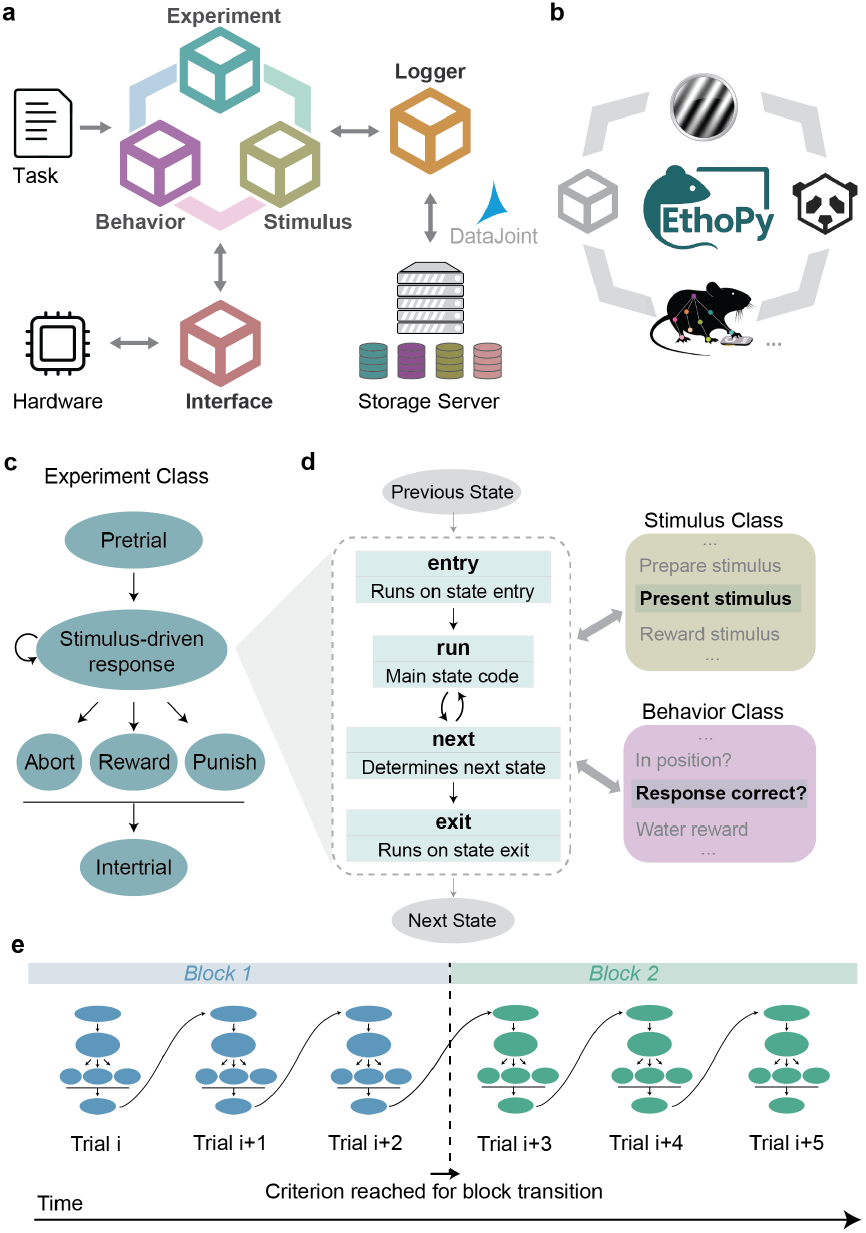
EthoPy software framework, showing the core modules, their interaction within EthoPy and with other relevant scientific communities. **a**. The task configuration file (Task) defines the parameters of the experiment (e.g., session start/end time, type of experiment, stimulus properties). The experiment module defines the experimental states (e.g., pretrial, punish period, etc.). The behavioral module handles the animal behavior (e.g., port selection, licking activity). Stimulus module creates and handles the presented stimuli. The Interface module handles the communication of EthoPy with the hardware (e.g., RPs, PCs). The Logger module handles the interaction with the storage server where data is saved using DataJoint. **b**. EthoPy is compatible and easily incorporates other publicly available and widely-used software packages, such as DeepLabCut, PsychoPy toolbox, and Panda3D. In addition, EthoPy modules are independent and thus can be easily modified and shared by the research community. **c**. In the Experiment class, a trial consists of multiple states and there are predefined rules for transitioning between states. Indicatively, the pretrial state is the period before the trial initiation, where the stimulus is prepared, and the intertrial state is the state following the end of a trial. Within a trial, a response based on the presented stimulus drives the transition to the outcome states (abort, reward or punish). **d**. A state is defined by 4 overridable methods, the “entry”, “run”, “next”, and “exit” functions. In each of these methods, a set of actions takes place using the methods of Stimulus and Behavior classes. For example, the stimulus-driven response state uses the Stimulus class for stimulus presentation in the “run” method and the Behavioral class for the animal’s response in the “next” method, to proceed to the outcome states. **e**. A trial is defined by specific experiment, stimulus, and behavior conditions. Trials can be grouped together to create a block, with a criterion for transitioning between blocks during a session.

Each module’s hyperparameters and data are stored into distinct but linked tables in a database server via Datajoint, the quickly expanding operating platform for scientific data management (Yatsenko et al., 2018; Johnson et al., 2024), maintaining full experimental traceability, including parameter changes and code versions. The database server further enables fast, combinatorial queries across large datasets and supports concurrent and distributed access for data sharing. Database’s dynamic management and balancing of synchronous data streams, guarantees scalable acquisition across many concurrent setups (Fig. 2a). Importantly, EthoPy achieves millisecond level precision (Supp. Fig. 1), by distributing time sensitive computations across processes and threads, supporting real time operations even on resource constrained platforms like the Raspberry Pi (RP).

**Fig. 2.**
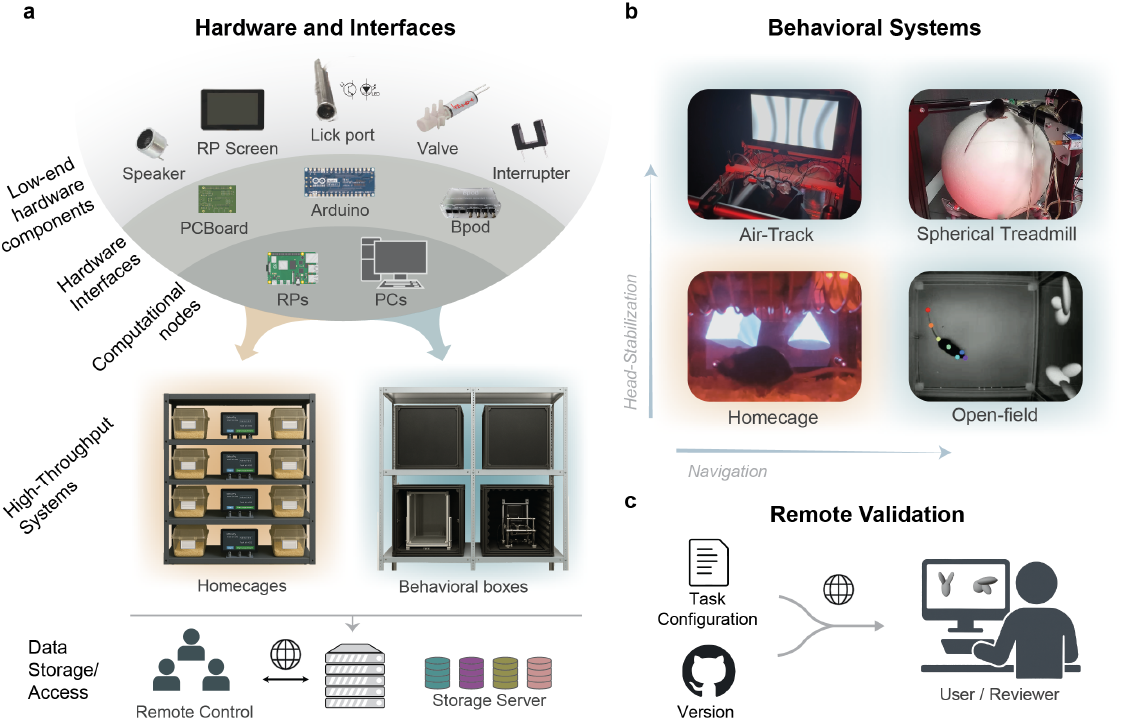
EthoPy supports diverse hardware implementations. **a**. Off-the-shelf or custom made low-end components, communicate via hardware interfaces, such as Arduino with custom PCBoard or Bpod, with the computational nodes that run the EthoPy (RPs, PCs). By using typical homecages modified only with 3 small openings, we can perform the behavioral experiments at the racks where the animals are housed. EthoPy also supports various different behavioral systems in larger behavioral boxes. Through the online communication with the storage server, users can remotely control experiments. **b**. Exemplar behavioral systems supported by EthoPy. Homecage: an RP behavioral setup, including a screen, is placed in front of a modified homecage with 3 holes for the ports. Air-Track: Minor modifications to the Homecage system allow for performance of the same behavior under head-fixation, using the Air-Track system. Openfield: Tracking of the animal’s pose through real-time video analysis is integrated in the EthoPy framework. Spherical Treadmill: Navigation under head-fixation using a spherical treadmill. Tracking of the animal’s position is performed by two sensors that encode the sphere’s movement. Axes indicate head-stabilization and navigation properties to account for diverse experimental needs. Background colors indicate development in homecage or behavioral boxes. See also Supp. Movie 1. **c**. EthoPy’s remote testing and validation of behavioral task configurations and precise version control enable collaborative development.

EthoPy promotes its adaptation from behavioral labs by leveraging Python’s widely-used and interpretable language (Muller et al., 2015) to support adoption in behavioral labs. Its high-level syntax simplifies extension, as most key tools (i.e., panda3d (Goslin and Mine, 2004), DeepLabCut (Mathis et al., 2018; Kane et al., 2020), PsychoPy (Peirce, 2007, 2008), OpenEphys (Siegle et al., 2017), etc.) are Python-based (Fig. 1b). EthoPy also exploits Python’s ecosystem to generate dynamic parameter sets, replacing static configuration files, and uses file-based logging for debugging and monitoring, all with negligible performance cost on modern processors.

#### Hierarchical architecture

The framework is organized into five core modules: experimental control (Experiment), behavioral monitoring (Behavior), stimulus presentation (Stimulus), hardware communication (Interface), and data management (Logger) (Fig. 1a). Each module implements core functions via abstract base classes that are further specialized through user-defined plugins to support specific hardware or workflows (see Methods section: EthoPy Plugins). This inheritance-based design allows integration of custom modules without rewriting core logic. Experiments are then configured through a task file (Task, Fig. 1a) that specifies session parameters and selected classes. This approach creates a two-tier customization model in which standard experiments require only parameter adjustment, while novel protocols can be developed through custom classes.

#### Experiment module constitutes the central control system of EthoPy

Experimental protocols are implemented as object-oriented state machines (Gamma et al., 1994) where states define specific behavior–stimulus interactions and transition rules (Fig. 1c). Each state is implemented as an independent class, which avoids the nested, conditional experimental logic of centralized approaches. Each state includes methods for “entry” (executed upon entering), “run” (main computational loop), “next” (transition logic), and “exit” (executed upon leaving) (Fig. 1d). The above architecture supports states’ reuse when implementing similar behavior–stimulus interactions (e.g., reward or punish). States automatically share their variables, allowing continuous tracking of trial history and transfer of experimental information between them; this supports complex experimental protocols that require coordination between different experimental phases. A trial is composed of multiple states and is defined by experiment, behavior, and stimulus parameters (Fig. 1c, Supp. Fig. 2, 3, 4). The methodology for trial selection can be tailored to the specific experimental needs, ranging from pseudorandomized selection to bias mitigation strategies. Sets of trials form blocks, with transitions determined by user-defined criteria such as accuracy, d-prime, or trial number (Fig. 1e). This architecture supports a wide range of implementations, from basic procedures, such as cleaning and port calibration, to monitored water delivery and complex tasks, like the 2 alternative forced choice (2AFC).

#### Behavior module forms the foundation for interpreting animal interactions

The Behavior module processes behavioral events such as position, proximity to target, and licking, while also tracking their duration and history. These events are acquired through the Interface module, and their interpretation within this module preserves a clear separation between event detection and behavioral logic. The Behavior module further validates behavioral conditions by evaluating their completeness for consistent data storage (Supp. Fig. 3). By extending this core functionality via inheritance-based classes, the Behavior module supports diverse paradigms, from simple response detection to spatial navigation tasks requiring dynamic position tracking.

#### Stimulus module handles stimuli presentation

The Stimu-lus module defines the full lifecycle of a stimulus, from preparation and presentation to cleanup, and manages the stimulus conditions (Supp. Fig. 4). Stimuli can be pre-generated at the start of a session for efficiency (e.g., movie clips) or rendered in real time for flexible, non-standardized presentations, such as those that depend on the animal’s position. This approach makes it straightforward for researchers to develop and integrate new stimulus types within a standardized framework. Specialized implementations extend across visual, olfactory, and auditory modalities, as well as their combinations. The module also provides standardized tools for stimulus timing and hardware synchronization with recording equipment.

#### Logger module manages the data

The Logger manages event timing and data storage during experiments using a producer–consumer architecture (Tanenbaum and Bos, 2015) with a priority-based message queue that saves critical data first. Asynchronous processing separates data generation from storage, safeguarding time-critical functions. A dedicated worker thread maintains data integrity through error handling and retry mechanisms (e.g., duplicate detection and database connection recovery), while another thread updates the Control table with real-time metrics (current state, trial count, total reward, Supp. Fig. 5). The Logger also records session metadata, including software version, hardware configuration, and task file, maintaining reproducibility. The direct integration with the database through Datajoint, provides for real-time remote monitoring and control of distributed setups, as well as reliable data capture for post hoc analysis (Fig. 2a; Supp. Fig. 5, 6). EthoPy, via Datajoint (Yatsenko et al., 2018), provides database creation and data management directly in Python, eliminating the need for SQL expertise. It also supports export of experimental data to the Neurodata Without Borders (NWB) HDF5 format (Teeters et al., 2015; Rübel et al., 2022), for standardized formatting and community tool compatibility, to support broader research collaboration.

#### Interface module controls communication with hardware

The Interface base class defines standard hardware functions such as liquid delivery, response detection, stimulation, and device synchronization, supporting flexible setups built from low-cost, custom-made, or commercial components (e.g., photodetectors, photointerrupters, valves, Fig. 2a, Low-end hardware components). Specialized hardware Interfaces (e.g., Arduino with our custom PCBoard or the commonly used Bpod, Fig. 2a, Hardware Interfaces) link these components to PCs with different operating systems or RPis (Fig. 2a, Computational nodes). Specialized Interface classes running on the computational nodes, handle device control (e.g., liquid delivery) and sensor-signal detection (e.g., lick detection or motion tracking). Because computational nodes operate independently and are remotely managed (Supp. Fig. 5), multiple setups can run in parallel in low-cost homecages (Supp. Table 1) or behavioral boxes (Fig. 2a, High-Throughput systems and Data Storage/Access). For each session, users can define the hardware configuration of a setup, e.g., the number and type of ports, and these parameters are stored in the server to safeguard consistency and reproducibility (Supp. Fig. 7).

#### Behavioral system support

Based on this framework, we developed several automated behavioral systems (Fig. 2b, Supp. Movie 1): (a) a Homecage System for low-stress training of freely moving mice with minimal experimenter input. This is used with standard rodent cages that fit typical facility racks so animals are trained in a familiar environment; (b) an Open-field System for freely moving animals, integrating real-time pose tracking with DeepLabCut-Live (Kane et al., 2020) to study exploration and navigation during decision-making; (c) an Air-track (Nashaat et al., 2016) System with hardware adapted to be similar to the Homecage System; and (d) a Spherical Treadmill System that allows animals to navigate while being head-fixed. The Air-track and Spherical Treadmill systems support experiments requiring head stabilization, a prerequisite for many neural recording techniques. These examples highlight EthoPy’s flexibility across diverse experimental paradigms, achieved by separating experimental logic from hardware implementation. Namely, the same experiment can be executed with different Interfaces, independent of hardware configuration. This design also supports collaborative development by enabling task design and validation without physical hardware, using mock interfaces (Fig. 2c).

### Online monitoring of a behavioral session

During a session, the Logger records all events in real time, enabling live tracking of behavioral metrics such as responses, block transitions, and lick counts per port (Fig. 3a–c). Experiments can be monitored remotely via the storage server, by using either Jupyter notebooks or a dedicated web interface for live plotting and control (see Methods: Software Experimental Control and Monitoring). The control table tracks animals, experiment status, session progress (e.g., consumed liquid; Supp. Fig. 5), and detailed activity (Supp. Fig. 6). Post-hoc, the database enables precise queries across sessions, allowing variables such as trial outcomes or aborts to be tracked for task evaluation and refinement (Fig. 3d–f). To support both online monitoring and offline analysis, we provide a Python package for data analysis and visualization, including example Jupyter notebooks (see Methods: Software - Behavioral Data Analysis).

**Fig. 3.**
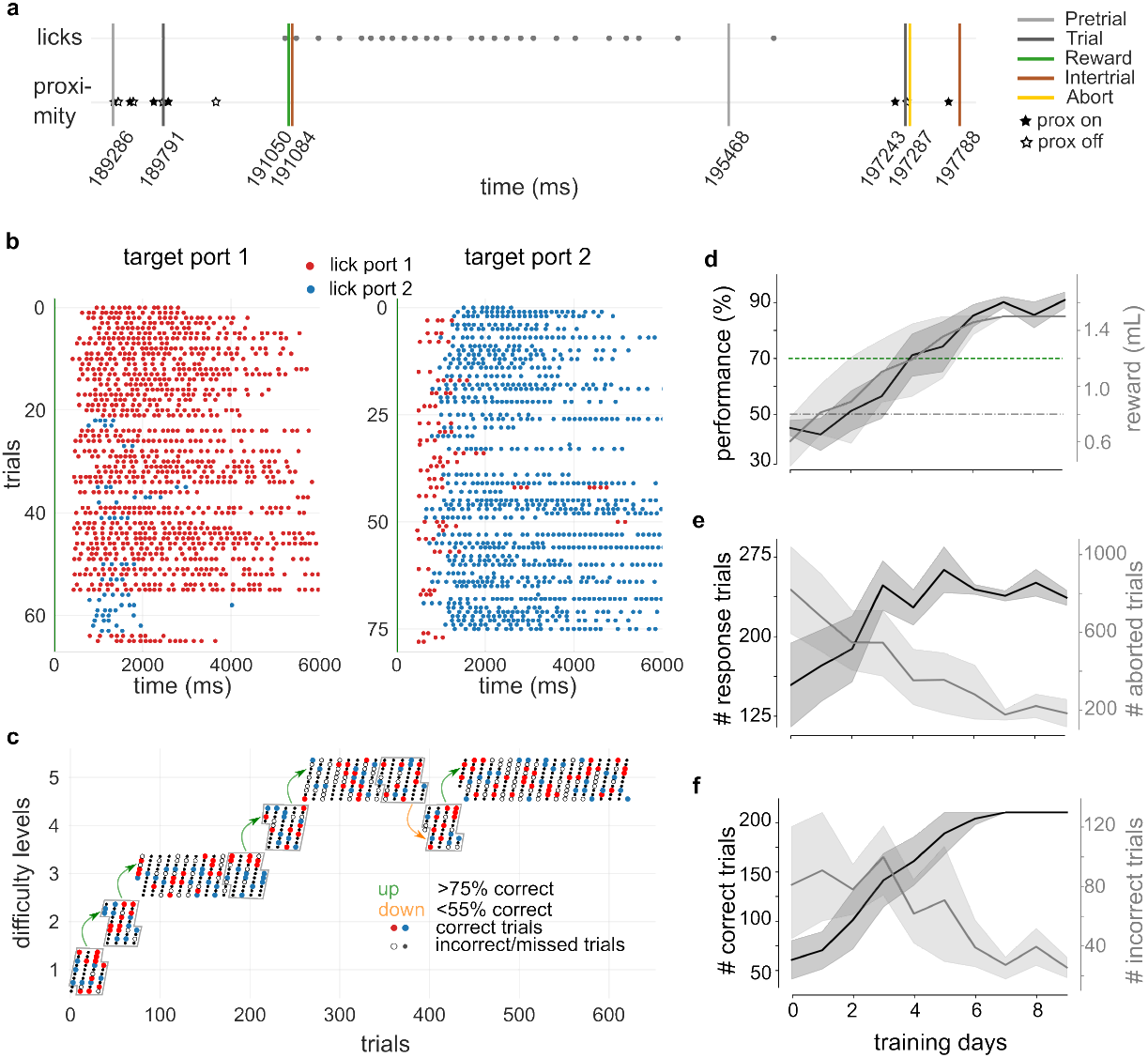
Behavioral monitoring during and across sessions. **a**. Timeline of two consecutive trials of a single session for a 2AFC task using 3D objects, that shows the distinct states (Pretrial, Trial, Reward, Intertrial, Abort) and animal responses (licks for receiving reward, and proximity on and off, i.e., nosepoke to initiate a trial). Time is in msec since the beginning of the session. **b**. Raster plot of licking activity following trial initiation (time = 0s, green) for a subset of trials for the same example session as in (a). Each dot is a lick and data are grouped by targeted rewarded port (left: target port 1, right: target port 2). Colors indicate licking to port 1 (red) or port 2 (blue). Note the extensive licking when the animal has made a correct choice, indicative of the availability of reward. **c**. The same session as in (a,b) showing block transitions: the mouse alternates between different blocks with increased task difficulty. The animal can transition to the next block or fall back to a previous block if its performance (evaluated in a moving 20 trials window, abort trials are not considered) is higher or lower than a criterion, respectively. Red (blue) dots correspond to trials that the correct port was port 1 (port 2) and the animal responded correctly; empty grey dots correspond to trials that the animal responded incorrectly, irrespective of port identity; small grey dots correspond to aborted or missed trials. **d**. Performance (black) and liquid consumption (grey) across training sessions of a simple object detection task, **e**. number of trials that the animal responded (either correct or incorrect, black) and number of aborted trials (grey) across training sessions, **f**. Number of correct (black) and incorrect (grey) trials across training sessions, for 8 mice performing a visual object detection task. Shading represents SEM.

### Diverse types of behavioral paradigms

We used the behavioral systems shown in Fig. 2b to implement several behavioral tasks that vary in complexity, from single-modality stimulus presentation and basic port selection, to multisensory environments and ethologically relevant behaviors involving active navigation. Specifically, with the Homecage system, we trained mice in different versions of the Go/no-Go and the 2AFC tasks, two widely used paradigms for probing sensory and cognitive functions in freely moving or head-restrained animals (Zoccolan and Di Filippo, 2018). Go/No-Go tasks are typically acquired rapidly, with high performance often reached in under five days for olfactory (Abraham et al., 2012) and auditory tasks (Ceballo et al., 2019). Indeed, in our Go/No-Go olfactory task (Fig. 4a, top), where mice were trained to discriminate between benzaldehyde (Go stimulus) and heptanone (No-Go stimulus), animals achieved high performance after only two sessions (Fig. 4a, bottom).

**Fig. 4.**
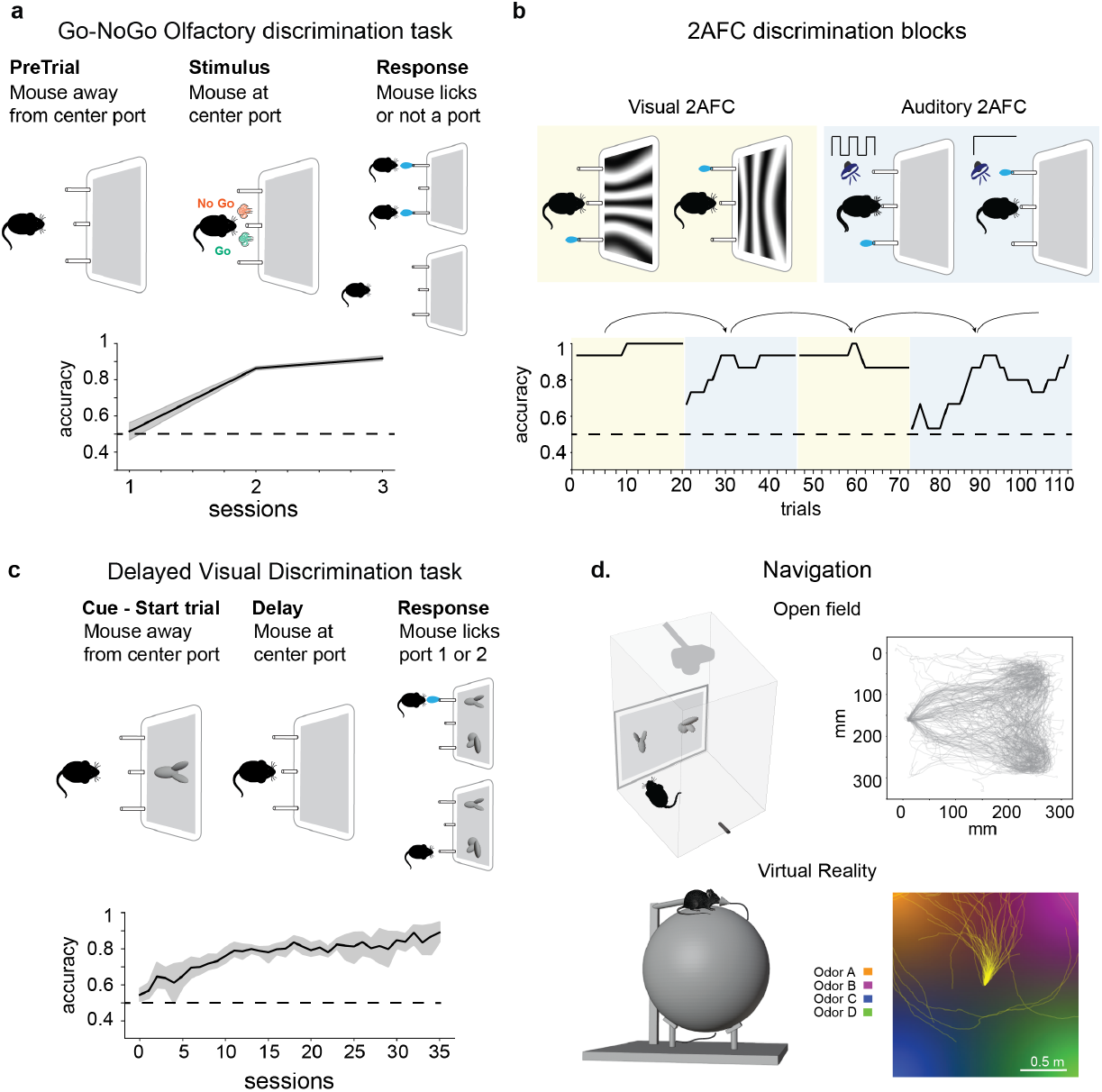
Diverse implementations of behavioral tasks using the EthoPy framework. **a**. Go/no-Go olfactory discrimination task (n=2 animals). The Go odor was benzaldehyde and the no-Go odor was heptanone. Once mice approached the center port (trial initiation), an odor was presented, based on which mice had to decide to lick or withhold licking to either port. **b**. Multiple block transitions within a single session of a mouse trained in two 2AFC discrimination tasks. Mice were trained to expert (>0.8) performance in a task requiring to discriminate between a 0 deg and 90 deg grating (Visual 2AFC), and in a task requiring to discriminate between 100 Hz tone clicks and a steady tone (Auditory 2AFC). Experimental states of a trial are the same as in a. Following training, mice alternated between the two rules multiple times within a session, with block switching induced once mice reached >0.8. Plotted performance is calculated with a moving average window size of 15 trials. **c**. Delayed Visual Discrimination task (n=5 animals). Different states can be implemented in EthoPy; here is shown the addition of the cue and delay period. During the cue period, a moving 3D object was presented. Following the delay period, mice were required to select the port on top of which was the cuestimulus. The port location of the distractor and cue stimuli varied between trials. **d**. Navigation tasks in the Open-field (top) and Virtual Reality using theSpherical treadmill (bottom) behavioral systems; (left) schematics of the behavioral systems, (right) trajectory lines during the trial period of an example session for each case. For the open-field visual discrimination task, the mouse had to ignore a distractor and approach the target stimulus to receive reward. Positioning of the target stimulus varied between trials. For the virtual reality olfactory discrimination task, the mouse could identify the direction of the target odor (Odor A) and move towards that corner to receive reward. For a-c dashed lines indicate chance performance.

2AFC tasks increase complexity by requiring animals to choose between two alternatives, with a response mandated on every trial to control for fluctuations in engagement (Zoccolan and Di Filippo, 2018; Dudchenko, 2004).

They have been applied across sensory modalities, including vision (Gibson and Maunsell, 1997; Froudarakis et al., 2020), olfaction (Roddick et al., 2014), and audition (Sakurai, 1990), but typically require longer training than Go/No-Go tasks. To address this, we used EthoPy’s block structure (Fig. 1e) to implement adaptive learning protocols, where animals advanced to harder blocks upon reaching performance criteria or reverted to easier ones when performance declined (Fig. 3c). Blocks also support studies of task switching in studies of decision making (Sul et al., 2011; Findling et al., 2023). As shown in Fig. 4b, mice performed multiple block transitions within a single session between two previously-learned visual and auditory discrimination tasks (see Methods: Behavioral Tasks for details). Formalizing protocols as sequences of discrete states (Fig. 1c) enables extending 2AFC tasks with distinct trial epochs, as in Match-to-Sample designs. These tasks probe cognitive processes such as memory, stimulus categorization, and decision making under specific manipulations and have been adapted to multiple different modalities (Gibson and Maunsell, 1997; Roddick et al., 2014; Sakurai, 1990). We implemented a visual delayed discrimination task (Fig. 4c, top) in which animals viewed a 3D moving object during the cue period, then identified the target among two comparison stimuli during the response period. Mice reliably discriminated between objects after 7–10 days of training (Fig. 4c, bottom).

Many ethologically relevant behaviors depend on locomo-tion and active sensing to reach a goal. Navigation inherently involves movement through space, whether for exploration, foraging, or goal-directed actions. To incorporate locomotion in freely moving animals, we developed an Open-field 2AFC visual discrimination task by integrating DeepLabCut-Live (Kane et al., 2020) into EthoPy for realtime choice inference based on position (Fig. 4d, top). In this setup, a screen displaying two 3D objects was placed opposite a reward port. Animals selected the target stimulus by staying within a predefined zone in front of it and then moved to the reward port. Trials were initiated only when the animals were away from the screen, preventing accidental responses to the presented stimuli. Training in this task induced selective movement between stimulus locations and the reward port (Fig. 4d, top right). To study navigation under head fixation, that is critical for investigating processes such as sensory processing (Niell and Stryker, 2010; Saleem et al., 2013), and spatial navigation (Dombeck et al., 2010; Domnisoru et al., 2013), we developed a virtual reality environment in EthoPy where mice navigated a spherical treadmill guided by odor cues (Fig. 4d, bottom). The environment was modeled as a square arena, each corner defined by a distinct odor whose intensity peaked locally and decayed radially. At any point, mice experienced a mixture of odors weighted by proximity, enabling spatial localization through relative intensity cues. By assessing head-fixed mouse trajectories (Fig. 4d, bottom-right) we observed that the continuous mixture of odors enabled spatial localization through relative intensity cues.

Together, these applications highlight EthoPy’s versatility across stimuli (visual, olfactory, auditory), interfaces (Raspberry Pi, Arduino, DeepLabCut, cameras, treadmill), behaviors (port selection, open-field, virtual reality navigation), and paradigms (Go/No-Go, 2AFC), while integrating community tools in both freely moving and head-restrained mice.

## Discussion

EthoPy offers a robust and customizable framework for training animals in a wide array of behavioral tasks using rich stimuli, making it a valuable tool for addressing diverse challenges in neuroscience research. Developed in Python (Muller et al., 2015), a widely adopted language in scientific computing for its versatility and cross-platform compatibility, EthoPy is accessible to users with varying levels of programming expertise. This design also enables rapid customization and extension to meet diverse experimental needs. As an open-source framework, EthoPy supports community-driven development through its modular architecture and plugin system, allowing researchers to contribute new components and adapt the platform to emerging paradigms. This fosters long-term sustainability and collaborative innovation across the neuroscience community (Gau et al., 2021; Diamantaki and Papoutsi, 2024). By leveraging off-the-shelf, low-cost components, EthoPy makes it feasible for laboratories to deploy sophisticated behavioral training setups at scale without the financial or logistic burden of proprietary systems. This affordability (Supp. Table 1) is particularly advantageous for labs aiming to train large cohorts or run high-throughput experiments in parallel. It also enables the execution of mock experiments without requiring specialized hardware, allowing researchers to validate or refine experimental protocols with ease before deploying them in full experimental settings. EthoPy’s flexible architecture further allows its adaptation not only to established behavioral paradigms such as the Morris water maze or Y maze (Dudchenko, 2004; Vorhees and Williams, 2006), but also to more complex visual navigational tasks using Panda3D (Goslin and Mine, 2004), further broadening its applicability across neuroscience research.

A distinguishing aspect of EthoPy is its integrated data management system, which uses the open-source Data-Joint framework to automate the organization, storage, and retrieval of experimental data and metadata (Yatsenko et al., 2018; Johnson et al., 2024). By embedding structured data management into its core architecture, EthoPy puts the FAIR principles (Martone, 2024) into practice, ensuring that experimental data and metadata are systematically captured, centrally stored, and readily accessible. Its integration with a relational database enables consistent logging of every parameter, from task configurations to software versions, facilitating traceability, version control, and reproducibility. In addition, support for exporting data in the NWB format (Teeters et al., 2015; Rübel et al., 2022) promotes standardized and transparent data sharing among research groups. Together, these features strengthen the long-term usability and scientific value of behavioral datasets, advancing the broader goals of open, collaborative neuroscience.

EthoPy’s automated nature significantly reduces the need for direct interaction between the experimenter and the animals, which is crucial for maintaining stable experimental conditions and minimizing behavioral variability (Richter, 2020; Nigri et al., 2022). This reduction in experimenter involvement is particularly beneficial in high-throughput scenarios, where the system can efficiently train large numbers of animals on complex tasks with minimal effort. Moreover, one of the most innovative aspects of EthoPy is its ability to facilitate training within the animals’ homecages with adaptive learning protocols that optimize training efficiency and promote a stress-free learning environment (Chesler et al., 2002; Sorge et al., 2014; Balcombe et al., 2004). This approach allows animals to learn challenging behavioral tasks without the added stress of frequent handling or changes in environment, which can often lead to inconsistent performance. Once trained, animals can be moved to more demanding tasks, such as those requiring head immobilization for neural recordings, with a solid foundation of learned behavior. This continuity between stress-free training and advanced experimental setups not only improves the reliability of behavioral data and significantly reduces experimenter workload, but also enhances the overall welfare of the animals involved. Importantly, concerns about stress from social isolation in homecage training were not evident in our experiments. The rapid learning rates we observed (Fig. 4a,c) suggest that automated schedules and minimal experimenter intervention help mitigate potential effects, aligning with reports that single- and socially housed animals show similar motivation and learning (Poddar et al., 2013). While laboratory based behavioral paradigms inherently involve artificial constraints that diverge from natural ethological contexts, EthoPy’s design philosophy prioritizes incorporating naturalistic elements wherever experimentally feasible.

Overall, by combining accessibility, scalability, modularity, and data integrity, EthoPy represents a significant step forward in behavioral training systems. It enables rigorous, high-throughput behavioral neuroscience and it is designed for extensibility, anticipating the rapid evolution of experimental needs across animal models. Looking ahead, its flexible design and strong community-oriented foundation make it well-suited for continued development, whether to support new hardware, integrate neural feedback, or promote standardized behavioral protocols across research groups.

## ACKNOWLEDGEMENTS

We would like to acknowledge Dr. Mostafa Nashaat for contributing with the Air-track behavioral system, and the members of the Froudarakis Lab for discussion and constructive feedback on the manuscript. E.F. acknowledges support from a European Research Council (ERC) grant (ERC-2022-STG, NEURACT, Grant agreement No: 101076710), the Hellenic Foundation for Research and Innovation (HFRI) under the 2nd Call for HFRI Research Projects to Support Faculty Members and Researchers with Grant Agreement No. 4049, and the Hellenic Foundation for Research and Innovation (HFRI) under the “Funding of Basic Research (Horizontal support of all Sciences)” of the National Recovery and Resilience Plan “Greece 2.0” with funding from the European Union - NextGenerationEU with Grant Agreement No. 016552. M.D. acknowledges support from the European Union’s Horizon 2020 research and innovation program under the Marie Skłodowska-Curie Actions with Grant Agreement No. 101025482. A.P. acknowledges support from the FORTH-Synergy grant (FlexBe).

## AUTHOR CONTRIBUTIONS

Conceptualization, E.F.; methodology, M.D., A.E., A.P., and E.F.; formal analysis, M.D., A.E., K.G., A.P., and E.F.; investigation, all authors; resources, E.F.; writing – original draft, M.D., A.E., A.P., and E.F.; writing – review & editing, all authors; visualization, M.D., A.E., A.P., and E.F.; supervision, E.F.; project administration, M.D., A.P., and E.F.; funding acquisition, A.P., E.F.

## Materials and Methods

### Ethics and Animal Care

Both male and female C57BL/6J mice (Strain #: 000664, The Jackson Laboratory) and Thy1-GCaMP6sGP4.3 mice (Strain #: 024275, The Jackson Laboratory) were used for behavioral training.

At the start of the training, mice were at least two months old and weighed at least 20g. In total, 17 mice were used; 12 male and 3 female C57BL/6J, and 2 male Thy1-GCaMP6sGP4.3. Mice were single-housed under a 12-h inverted light/dark cycle and the experiments were conducted during the dark phase. Animals had unrestricted access to food and were maintained under water restriction, receiving between 800 µl and 1500 µl of water per day. Body weight was monitored regularly to ensure their health and well-being. All procedures were approved by the Veterinarian Authorities of the Region of Crete with protocol number 54506.

### Software

#### Code Availability & Installation

The EthoPy package is freely available at https://github.com/ef-lab/ethopy_package under the MIT License. Setup and installation instructions for both EthoPy and the associated database can be found in the EthoPy GitHub repository.

#### Database Schema Organization

The relational database used in EthoPy is managed through DataJoint (Yatsenko et al., 2018), which provides a high-level Python interface for interacting with SQL-based systems such as MariaDB and MySQL. The database is organized into schemas, i.e., named collections of related tables that organize the data. EthoPy defines separate schemas for the experiments, behavior, stimuli, interfaces, and recordings, each of which contains a subset of tables for the comprehensive saving of all experimental parameters and outcomes.

##### Experiment Schema

The experiment schema (Supp. Fig. 2) defines a structured relational database designed to manage the parameters and metadata associated with the experiment module. At its core, the schema organizes data around sessions, each representing a single experimental run for a specific animal (animal_id). The session table (Supp. Fig. 2, Session) stores metadata including the user conducting the session, the setup used, the experiment type, the session timestamp, the task configuration file, and the code versioning.

Each session is linked to its trials via shared primary keys (animal_id, session) (Supp. Fig. 2, Trials). Trial-specific information is stored in the trial table, and each trial is associated with its corresponding state transitions (e.g., Trial, Reward, Punish), which capture the temporal progression of states within a trial. Additionally, each trial references a unique set of experimental conditions (Supp. Fig. 2, Conditions), captured in the condition table and identified by a cond_hash. A hash is a unique computed identifier derived from the experimental parameters that acts like a digital fingerprint, ensuring that trials with identical conditions can be easily identified and grouped together. Conditions are further partitioned into specialized tables tailored to specific experimental paradigms. For example the default condition_passive, condition_match_port, and condition_free_water tables contain experiment-specific parameters such as reward amounts, intertrial intervals, and trial selection strategies. These sub-tables maintain relational integrity by using the same cond_hash as a foreign key to the condition table. Any new experiment should create a new sub-table as shown with the high opacity table called new_experiment (Supp. Fig. 2, where all the experiment parameters (conditions) will be saved.

The control table (Supp. Fig. 2, 5, Lookup Tables: control) provides real time operational metadata for each setup, including the current experiment status, active state, task index, trial count, reward delivery, and timing-related information. This table is critical for managing experimental execution by monitoring the current status of each setup. Finally, the task table (Supp. Fig. 2, Lookup Tables: task) serves as a lookup for task configuration files, linking a unique task identifier (task_idx) to a description, path, and timestamp of creation.

##### Behavior Schema

This schema (Supp. Fig. 3) is organized into two main sections: Activity and Conditions. Both sections are connected to the trial table (grey table) from the experiment schema (Supp. Fig. 2) via the shared primary keys animal_id, session, and trial_idx, ensuring that both behavioral events and behavioral metadata are tightly integrated with the trial structure. The Activity tables (activity, activity_position, activity_lick, activity_proximity) record behavioral events of the animal during each trial, including positional tracking, licking, and proximity to interaction ports. The Conditions tables capture the behavioral parameters assigned to each trial. Central to this is the beh_condition table, which is identified by a unique beh_hash and linked to condition specific parameter tables (e.g., multi_port, multi_port_reward, and multi_port_response) that define the target behavior, such as the correct response port for reward.

##### Stimulus Schema

The stimulus schema defines all conditions related to the creation and presentation of stimuli during experiments (Supp. Fig. 4). Each trial is associated with a unique stim_hash, which encapsulates the specific parameters of the stimulus used in this trial. The stim_condition table stores these parameters, while the stim_condition_trial table links them to individual trials and records the timing of stimulus presentation. Detailed configurations for each stimulus type (e.g., dot, bar, grating, grating_movie) are stored in dedicated subtype tables, all connected via the shared stim_hash.

##### Interface Schema

The interface schema manages all parameters related to the experimental hardware configuration. It is structured around two main components: setup configuration and calibration (Supp. Fig. 7). The setup configuration allows users to define the specific hardware components (e.g., ports, sensors, screen) to be used in an experiment by referencing predefined lookup tables. This design enables users to configure the entire setup within a task simply by specifying the setup configuration id, which in turn maps to all associated subcomponents and their parameters. To safeguard reproducibility, the session-specific setup configuration is stored separately for each session, capturing the exact hardware parameters (or lack of in the case of the mock interface) used at the time of the experiment, even if the lookup tables are later modified. In addition, calibration tables store important measurements of the hardware that might vary over time, such as water dispensation volumes for individual ports.

#### Task setup and execution

Tasks in EthoPy are Python scripts that define the experimental protocol through a combination of experiment, behavior, and stimulus parameters. Each task is structured with three main components: (1) import of the experiment, stimulus, and behavior modules needed for the particular experiment, (2) session parameters that control global experiment settings (e.g., setup configuration index, reward limits), and (3) stimulus, behavior, and experiment trial-specific conditions. Tasks can be created using the `ethopy-create-task` command line tool, which generates a template file containing all necessary parameters and placeholders. The template is then customized by specifying the required module paths and class names for the experiment, behavior, and stimulus components. Trial conditions are defined using the experiment’s Block class, which implements the staircase or other trial selection methods. The experiment is executed by pushing the defined conditions to the experiment object and initiating the start method.

A basic task implementation follows this structure:

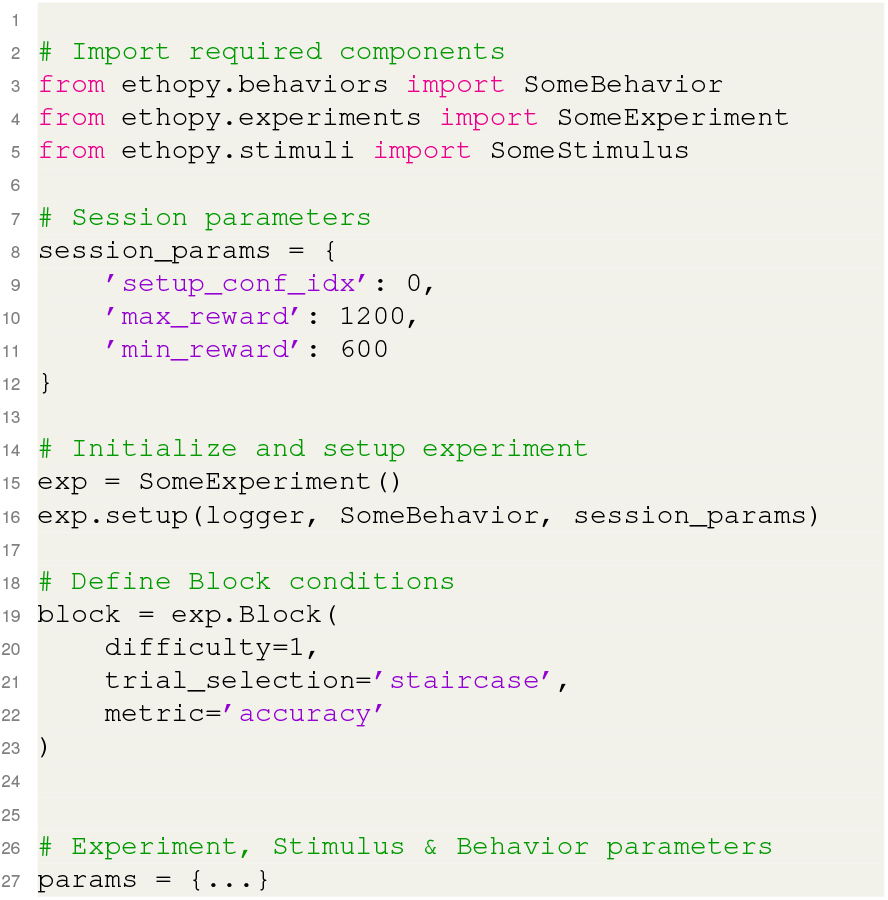
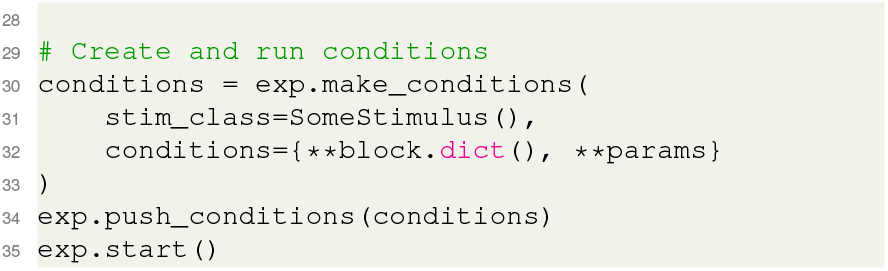

Tasks can be executed via command line with:

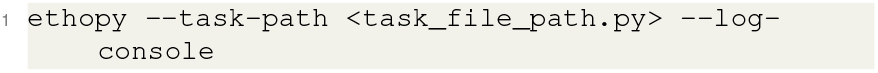

The task configuration file can be stored in the Task table (Supp. Fig. 2) with a unique task_idx, which serves as a numerical identifier that links the task configuration path to an id. The task_idx can be used in the Control table to specify which experimental protocol should run on a particular setup. The users simply update the task_idx field in the Control table to remotely trigger the execution of the corresponding task configuration (Supp. Fig. 2). Task can also be executed by specifying the task_idx via the command line:

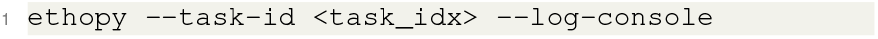

#### EthoPy Plugins

The EthoPy framework incorporates a flexible plugin system that enables the extension of core functionality through custom modules specifying behaviors, experiments, interfaces, and stimuli. Plugins can be implemented either as standalone, single module, or categorized components within specific directories (behavior, experiment, interface, and stimulus modules). The system automatically discovers and registers plugins both from default locations (`∼/.ethopy/ethopy_plugins/`) and custom paths specified through the `ETHOPY_PLUGIN_PATH` environment variable. Plugin resolution follows a hierarchical precedence where core EthoPy modules take priority over user defined plugins, with later added paths overriding earlier ones in case of conflicts. The plugin architecture allows for seamless integration of custom experimental protocols while maintaining compatibility with the core framework. Each plugin can be imported using the `ethopy` namespace (e.g., `ethopy.behaviors.custom_behavior`), enabling a standardized approach to extending experimental capabilities. The system includes built-in conflict resolution and management features through the `PluginManager` class, which handles plugin discovery, registration, and information tracking. This extensible architecture facilitates the implementation of specialized experimental paradigms while ensuring robust integration with the core EthoPy framework. Examples of plugins can be found at EthoPy Plugins GitHub Repository (https://github.com/ef-lab/ethopy_plugins/), providing a valuable resource for researchers looking to extend the framework.

#### Experimental Control and Monitoring

EthoPy Control (https://github.com/ef-lab/ethopy_control/) serves as a web-based graphical interface that complements the EthoPy behavioral experiment framework. While EthoPy provides the core state control system for behavioral experiments on individual setups, ethopy_control enables laboratory-wide coordination and monitoring of multiple experimental systems. The Control monitoring page (Supp. Fig. 5) through the Control table at the Experiment schema (Supp. Fig. 2) provides realtime status monitoring of all laboratory setups, displaying critical parameters such as status(e.g. running, sleeping), user, ping, and experimental progress across multiple behavioral systems. Each setup can operate autonomously on a daily schedule based on configurable start and stop times defined within the individual setup parameters, enabling automated experiment execution without manual intervention. The control page serves as the primary interface for researchers for assigning tasks to specific setups, monitor session progress, and perform remote setup edits, including system reboot. Additionally, the platform integrates realtime data visualization capabilities that display live behavioral events (lick port activations, proximity sensor data) across configurable time windows, enabling immediate assessment of experimental progress and animal performance (Supp. Fig. 6). A dedicated task management page enables researchers to create, modify, and organize task configurations by adding new task_idx entries (Supp. Fig. 8).

#### Behavioral Data Analysis

EthoPy Analysis (https://github.com/ef-lab/ethopy_analysis/) is a Python package that extends the EthoPy framework for behavioral data analysis and visualization. Built with a modular, DataFrame based architecture, this package provides DataJoint based tools for analyzing multi-session behavioral experiments across animals and sessions. The package abstracts database complexity by providing high-level data loading functions that retrieve behavioral data from DataJoint schemas and return them in pandas DataFrame format, eliminating the need for users to write SQL queries or understand DataJoint syntax. Key features include automated performance metrics calculation, standardized visualization functions for learning curves and behavioral patterns (Fig. 3), and a command-line interface for batch processing experimental datasets. This extension enhances the EthoPy ecosystem by providing analysis tools that support reproducible behavioral neuroscience research workflows.

#### Hardware

All the information mentioned below, with assembly instructions, parts list, and 3D designs, is in the EthoPy Hardware GitHub Repository (https://github.com/ef-lab/ethopy_hardware/).

#### Hardware components

##### Interruptor ports

Interruptor ports [HR0172] are mainly used as proximity detectors, i.e. detection when and for how long a subject is located near to the port. Interruptor ports include an aligned LED and photo sensor. The photo sensor detects the animal’s movement when it approaches the port through the infrared interruption. The photo sensor of the port is connected to the Hardware Interface (Fig. 2a) which links the port with the computational node (RP or PC).

##### Lick ports

Lick Ports can be used both for lick detection and reward delivery. Regarding the former, lick ports include an IR LED and a photodiode positioned in parallel. Close vicinity of an object (e.g., the tongue of a mouse) reflects the infrared light and increases the signal from the photodiode. The photodiode is connected to the control board that links it with the computational node (RP or PC). Regarding the latter, a tube for water delivery for correct responses is coupled to a computerized valve that controls the liquid delivery, which can deliver liquid volumes with 1µL resolution. The water is channeled through a liquid delivery device when the correct electrical signal is transmitted from the port to the electric control panel, leading to the opening of the respective valve. Both the LED and photodiode wires and the reward tubes are enclosed in a stainless steel 6mm diameter cylinder, and epoxy glue is used at their endpoints in order to prevent possible damage that could be caused by chewing or other mouse activities. In addition to detecting licks, the infrared-based sensing mechanism of the lick ports can also function as a proximity detector, allowing them to serve as a substitute for interruptor ports in certain experimental configurations (https://github.com/ef-lab/ethopy_hardware/blob/main/Homecage/Lick_ports_assemply.md).

##### EthoPy Controller Board and Arduino

To sup-port the behavioral system described in this study, we developed a custom circuit board (PCBoard) that serves as an interface hub between the low-end hardware components and the computational nodes (EthoPy Controller https://github.com/ef-lab/ethopy_hardware/tree/main/EthoPy_Controller).

This board connects two lick ports and a central trial initiation port to a signal-processing microcontroller and a reward delivery system. Specifically, the PCBoard connects with an Arduino Nano, which receives and processes input signals from the photodiodes of the lick and center ports, allowing precise detection of animal interactions (https://github.com/ef-lab/ethopy_package/tree/main/src/ethopy/interfaces/arduino_firmware). The PCBoard further incorporates the necessary electrical components to drive solenoid valves, enabling precise and reliable control of liquid reward delivery. It also features three onboard buttons: two for manually activating the solenoid valves to verify reward delivery, and a third used to calibrate the activation thresholds of the lick and center ports. A dedicated connector on the board provides seamless integration with the GPIO ports of an RP that runs the EthoPy framework. Alternatively, the Arduino can be connected directly via USB to a PC that runs EthoPy, providing additional flexibility in experimental setups. This compact and modular hardware architecture reduces wiring complexity and supports rapid assembly and scalability across multiple experimental setups.

##### RP behavioral setup

For this setup, an RP computer with an SD card with sufficient storage (recommended at least 32GB) is connected to a 7-inch RP display screen and the control board. The latter ensures the communication between the ports, the miniature solenoid valves, and the processing node, in this case the RP. The number and type of ports are set up according to the needs of the specific experiment. For example, for the setup shown in Fig. 2b, a proximity port - that can be either an interruptor or a lick port - is located in the center of the apparatus (‘center port’) and can be used for trial initiation. Two lick ports, on either side of the center port, are used to detect licks and deliver rewards. The subject’s response is recorded, timestamped and uploaded to the database, immediately available for visualization.

Using development boards like RP and Arduino makes the behavioral system much easier to work with. These standard platforms allow researchers to easily change and adapt their hardware. The system becomes less complicated to wire and set up, while staying consistent across multiple experimental systems. Development boards can make it simpler to connect custom sensors and devices, making advanced behavioral experiments more accessible to researchers and easier to reproduce in different laboratories.

### Behavioral systems

#### Homecage

We developed a Homecage behavioral system built on the above-described RP setups (Fig. 2b, Homecage). Each unit consists of a standard mouse cage [1264C EUROSTANDARD TYPE II] modified with three openings (∼7.5 mm diameter for lick ports, ∼8.5 mm for speakers if included) on one side. These openings align with the RP behavioral setup, which includes ports, optional speakers, and a 7inch RP screen mounted ∼41 mm from the central proximity port. The setup can be mounted on custom 3D-printed stands (https://github.com/ef-lab/ethopy_hardware/tree/main/Homecage/3d_designs) that hold ports, speakers, and the screen. The homecages can be placed directly onto standard racks in the animal facility, allowing mice to be trained in their familiar environment. Each unit operates as an independent node, connected directly to the central data management system, enabling multiple parallel experiments with minimal experimenter involvement. This design supports scalable, high-throughput behavioral training while reducing animal stress and handling. Room requirements for the system include: (1) power supply, (2) air supply (e.g., small compressor for liquid delivery), and (3) network connectivity (Wi-Fi or Ethernet).

#### Air-track

Many neural recording techniques require head-fixation during behavioral tasks. To accommodate this, we adapted the RP behavioral setup to control the Air-Track platform developed by Nashaat et al. (Nasr et al., 2022; Nashaat et al., 2016, 2024) (Fig. 2b, Air-Track). This adaptation involved minimal hardware changes - mainly adjusting the photodetectors and the lick ports to fit the floating, air-cushioned platform. In the Air-Track configuration, an interruptor-style behavioral port (e.g., Bpod mouse behavior port https://sanworks.github.io/Bpod_Wiki/assembly/mouse-behavior-port-assembly/) aligns with platform requirements, and photodiode sensor signals are processed as in other EthoPy setups to trigger behavioral events. Thanks to EthoPy’s modular architecture, these modifications required no changes to the core RP-based control system. The RP behavioral setup can be securely mounted in front of the Air-Track using custom 3D-printed holders, maintaining precise stimulus control and response detection. This compatibility allows animals to be pre-trained in the homecage and then transition to head-fixed experiments.

#### Open-field

Studying naturalistic behaviors such as spatial exploration, decision-making, and goal-directed movement requires tasks that allow animals to navigate freely within their environment. To this end, we developed an open-field behavioral system (Fig. 2b, Open-field) for investigating exploration and navigation in freely moving mice. The system consists of a transparent Plexiglas arena (30 × 30 × 35 cm) enclosed within a sound and light isolated box [RatRig: https://ratrig.dozuki.com/Guide/01.+V-Hive+Enclosure+Base+Model/183?lang=en], providing a controlled testing environment. A 13.3-inch monitor is mounted on one side for visual stimulus presentation, with a single lick port on the opposite side for reward delivery. Both the port and reward delivery hardware are controlled via the same custom circuit board used in other setups via serial communication with a PC. An overhead camera, aided by red LED lighting for uniform, non-intrusive illumination, tracks the animal’s position in real time. Tracking is powered by the integration of DeepLabCut-Live (Kane et al., 2020) to EthoPy, enabling closed-loop task execution based on the animal’s position. Trials and state transitions (Fig. 1c) are triggered by location-dependent events - for example, starting a trial when the animal enters a defined area, or delivering reward/punishment based on its position relative to the stimulus. For real-time inference at >30 FPS, EthoPy runs on Ubuntu with an NVIDIA GPU (minimum 8 GB VRAM) to maintain TensorFlow compatibility and GPU-accelerated processing. Data from each session, including trial events, pose coordinates, and session videos, is uploaded immediately to the database for visualization and storage. At the end of the session, the locations of the video and a file with the animal coordinates are saved in the database.

#### Spherical treadmill

To investigate navigation-based behaviors under head-fixation, we developed a virtual reality (VR) behavioral system using a spherical treadmill that allows animals to navigate through simulated environments (Fig. 2b, Spherical treadmill). In this case, the RP behavioral setup incorporates two motion sensors for extracting translational movement, along with odor and reward delivery systems. The spherical treadmill setup consists of a 25cm diameter Styrofoam ball supported by three metal ball bearings, enabling smooth, low-friction movement. Animals are head-fixed via two metal posts on top of the ball, and navigate by rotating it. Movement is tracked using two optical computer mice placed 90°apart and positioned close to the bearings for accurate motion detection. Odor stimuli are delivered via a custom-built manifold that also includes a lick port for reward delivery. Licking is detected using a capacitance-based detection circuit [SEN-14520]. Odor delivery is managed by a custom olfactometer capable of switching between four odors through solenoid valves driven by PWM signals from an RP running EthoPy (see Olfactory stimuli). The setup requires the same infrastructure as other systems—power supply, air supply, and internet connectivity.

### Operational Procedures

#### Liquid delivery

In the animal facility room, a dedicated water delivery system is required to provide consistent and precise liquid rewards across multiple behavioral setups. The system consists of an air supply with adjustable pressure, an airtight container filled with potable water or other reward solution, a pressurized tubing network, and inline micropore filters (45µm pore size). The air supply, typically set between 3-8psi, is connected to the sealed water container to maintain constant pressure. This pressure forces water through the food-safe tubing system towards the behavioral setups, where solenoid valves control the release of liquid through the lick ports. Maintaining constant pressure facilitates fast, consistent water delivery with minimal latency. Micropore filters are placed near the end of the tubing system just before the valves to prevent clogging from particulates or debris. All tubing and fittings should be leak-proof and securely connected to avoid air loss or contamination, ensuring reliable performance during training sessions.

#### Cleaning

The water delivery system requires regular cleaning - at least once a week - to prevent contamination and clogging of tubes and lick ports. Cleaning is performed using a 0.7% detergent solution of Nalgene L900 Liquid Detergent and increased pressure set at 10psi. Each port is flushed with 70 pulses of 300ms duration of liquid, and 100ms intervals between pulses. The same procedure is repeated using purified water to rinse out any detergent residue. Twice a month, an additional disinfection cycle should be carried out using a 5% chlorine solution. Micropore filters (see above) must be removed before starting the cleaning procedure and replaced with new filters upon completion to maintain consistent flow across long time periods.

#### Calibration

To secure precise and consistent reward delivery across all behavioral setups, each liquid port is calibrated to determine the relationship between valve opening duration and the volume of water released. Calibration is performed weekly after cleaning the system and is typically carried out under a constant air pressure of ∼5psi. For each port, the solenoid valve opens for a series of predefined durations, typically 20ms, 35ms, and 100ms. For each duration, the valve is triggered multiple times to account for variability and obtain an average flow rate. The released water is collected and weighed using a precision scale. These measurements are then entered directly into the system via EthoPy’s graphical interface for each setup. EthoPy uses these measurements to compute the flow rate and generate a calibration curve, allowing it to automatically determine the exact valve opening duration required to dispense a specified reward volume.

#### Synchronization

To establish accurate timing and synchronization across behavioral, visual, and recording components, we characterized key sources of temporal delay and system latency in our experimental setup. During the behavioral task, the average delay between a lick response and the subsequent activation signal for the reward valve was estimated to be 1.2ms (Supp. Fig. 1a). To assess real-time rendering performance, we measured the interval between two consecutive frame flips while presenting moving visual stimuli rendered using Panda3D (Goslin and Mine, 2004) on an RP4. The monitor operated at its native refresh rate of 60Hz (Supp. Fig. 1b), demonstrating the system’s ability to maintain consistent frame timing. We also quantified the monitor’s display response latency using a photodetector, finding an average lag of 30ms between the frame flip command and the actual visual update (Supp. Fig. 1c). For cross-system synchronization — including behavioral, recording, and database server — we employed Precision Time Protocol (PTP). On RP4, which supports only software-based timestamping, the average synchronization precision was estimated at approximately 1.4ms (Supp. Fig. 1d). In contrast, systems supporting hardware-based PTP timestamping (such as RP5 or desktop machines with compatible network interfaces) can achieve sub-millisecond precision. EthoPy also supports syncing by generating/recording digital pulses across different devices.

### Stimuli

#### Olfactory

Odor delivery is controlled via a pulse-width modulation (PWM) signal — generated from the RP’s GPIOs — that drives a solenoid valve circuit. The resulting pulses produce brief “puffs” of air, which pass through bottles containing diluted odorants and are directed to the animal via the center port (Fig. 4a). Odor concentration can be algorithmically modulated by varying the ON/OFF timing of the PWM signal, or manually adjusted by diluting the odorant in a carrier solvent such as mineral oil. EthoPy also supports mixtures of odorants and dynamic switching between them. In the olfactory task shown in Fig. 4a, an aldehyde (50% benzaldehyde in mineral oil) served as the target odor gradient, while a ketone (10% heptanone in mineral oil) was used as the distractor during the “no-go” phase. For the VR navigation task (Fig. 4d, bottom), the 4 odors used are propyl-acetate, 2-heptanone, and benzaldehyde (50% in mineral oil) and carvone (not diluted).

#### Auditory

For tone delivery, the speakers are directly controlled from the output of the GPIOs of the RP. Modulation of this electrical signal can generate tone ‘clicks’, as the ones shown in Fig. 4b. In the homecage behavioral system, we used Ultrasonic Piezoelectric Transducers as speakers tuned at 40kHz; this was necessary as ultrasounds are characterized by a very focused spatial profile (high reduction in amplitude of the ultrasound beam as a function of distance) that allows for multiple auditory experiments to be performed within an animal room without interference between the cages.

#### Visual

##### Objects

For presenting 3D objects, EthoPy integrates Panda3D (Goslin and Mine, 2004), an open-source engine optimized for real-time visualizations. This enables precise, dynamic control over both the objects and their visual context during experiments. Parameters such as object size, position, rotation, and motion can be flexibly adjusted, along with background features including color, lighting direction, and intensity. In the tasks shown in Fig. 4c,d, we presented two moving 3D objects against a black or gray background, systematically varying their size, horizontal and vertical position, rotation, tilt, yaw, and illumination to explore the animals’ ability to discriminate complex visual features.

##### Gratings

PsychoPy (Peirce, 2007, 2008), a Python-based open-source package, has been integrated into EthoPy to enable the creation and manipulation of visual gratings. This integration allows for the flexible control of various visual parameters, including grating position along the x-and y-axes, masking, orientation, and spatial and temporal frequency, tailored to the needs of each experiment.

### Behavioral Tasks

#### Training procedure

For all behavioral systems (Homecage, Air-track, Open-field, and Spherical treadmill), the training procedure is divided into two main stages: (1) habituation to the setup and (2) training to the specific behavioral task.

In the case of the homecage behavioral system, during habituation, animals first learn to lick freely from lick port(s) to receive water, which helps them associate the ports with reward and become familiar with the setup. In a subsequent session, they are required to initiate each trial by activating the center port before gaining access to the side ports. This promotes familiarization with the trial structure. Habituation is typically completed after the animal manages to drink at least 1mL of water within 2 hours, and it usually takes one day (maximum 2 days) depending on the individual animal.

Habituation to the Air-track behavioral system, Open-field arena and Spherical treadmill requires transfer of the animal from the homecage to the setup. In the Air-track behavioral system, mice are initially freely moving and are habituated to the task structure similar to the homecage behavioral system (∼2-3 days). Once trial initiation is learned, mice are further habituated to head-fixation, requiring also to learn to move the air-platform (∼7 days), before training to a specific task. In the Open-field, animals are initially freely moving and learn to lick freely from the lick port to receive water (∼1-2 days). In the following days, mice are allowed to move freely for approximately 10–15min before task initiation to habituate and reduce stress following transfer from the homecage. Habituation to the spherical treadmill requires habituation to the headfixation (2 days) and to the lick port (∼2-3 days), where the reward odor is also introduced. During this phase, animals also have to learn how to walk on the spherical treadmill.

Once habituated, animals begin training on the specific behavioral tasks. Water is made available exclusively through task performance, with a daily minimum of 0.7mL and a maximum of 1.5mL. Food remains freely available in the homecage. Training sessions typically last 1–3h per day. All task parameters, including trial structure, stimulus timing, and reward delivery, are configurable based on experimental requirements. EthoPy supports several approaches for controlling task progression, including performance-adaptive paradigms. In the latter case, task difficulty automatically adjusts based on the subject’s performance (Treutwein, 1995; Garcia-Perez, 1998; Leek, 2001): if performance exceeds a threshold (e.g., 80% correct), the task becomes more difficult; if it falls below a minimum threshold (e.g., 50%), the difficulty is reduced (Fig. 3a). This adaptive training allows animals to progress within the same session.

#### Go-NoGo task

In a Go-NoGo task in the homecage setup (Fig. 4a), a trial begins when the mouse places its snout at the center port for a minimum of a predefined period (∼100ms). A stimulus is then presented in the form of an odor for 500ms. To facilitate engagement, the animal must maintain its position for an additional 100ms after stimulus presentation. Each trial is designated as either a “go” trial (target odor) or a “no-go” trial (distractor odor). During “go” trials, the mouse can receive water reward by licking at either lick port (correct response). If no licking response occurs within 3s, the trial is terminated and marked as aborted (miss). During “no-go” trials, the correct behavior is to withhold licking for 3s (correct rejection). If the animal licks during these trials, the response is marked as incorrect (false alarm) and results in a 10s timeout penalty (punish).

#### Two alternative forced choice tasks

In a standard 2AFC task in the homecage setup, as the ones shown in Fig 4b, a trial begins when the mouse places its snout at the center port and stays in place for a minimum of a predefined period (50ms). Then either a stimulus appears on the screen or a sound is activated, and the animal has 5s to make a response by approaching one of the two lick ports. Each stimulus is uniquely paired with a specific lick port. If the mouse licks the correct port, the screen turns white and a water reward (5-10µL) is delivered through the lick port. If the mouse licks the wrong port, the screen turns black, and there is a timeout penalty of 4-8s (punishment) before the next trial. If there is no response to either port within 5s, the trial is terminated and marked as aborted. For block switching of Fig. 4b, a new set of trials was presented once the animal reached >80% performance calculated in a 20-trial window.

In the cued 2AFC task (Delayed Visual Discrimination task) in the homecage setup shown in Fig. 4c, a trial starts automatically with the presentation of the target object in the center of the screen (Cue period). For the implementation of the moving objects we used Panda3D (Goslin and Mine, 2004). During the Cue period, the animal has time to sample (view) the target object. For the example data in Fig. 4c the Cue period was set to a maximum of 4min. After sufficient sampling of the target object, the animal must approach and maintain its snout at the center port for 0.2-0.5s (Delay period). Subsequently, the target object appears on one side of the screen, and a distractor object appears on the opposite side (Response period). During the Response period, the animal must identify the target object and lick the port in front of the target object. If the animal’s choice is correct, the screen turns white and a water reward (5-10µL) is delivered through the lick port. If the choice is incorrect, meaning the animal licked the port in front of the distractor object, the screen turns black, and there is a timeout penalty of 4-10s (punish) before the next trial.

#### Open-field Visual discrimination task

In the open-field visual discrimination task (Fig. 4d), the animal position is recognised and a trial begins when the animal is in a predefined area of the arena. Then, two stimuli appear on the screen (the target object on one side of the screen and the distractor object on the opposite side). The animal must respond by approaching one of the stimuli. If the mouse approaches the correct stimulus (target), the screen turns white and a water reward (5µL) is delivered through the lick port that is positioned on the opposite side of the arena. If the mouse approaches the wrong stimulus (distractor), the screen turns black, and there is a timeout penalty of 20s (punishment) before the next trial.

#### Virtual reality olfactory discrimination task

In the head-fixed VR navigation using an olfactory discrimination task, the animal has to approach the target odor to receive reward (∼5-8µL of water). The trial starts with the animal positioned in the center of the virtual space, where it receives an equal amount of all 4 odors. Then, the animal must move through the virtual space towards the higher concentration of the rewarding odor. The reward can be granted once the animal enters a pre-specified area centered at the source of the rewarding odor. The reward is delivered only if the animal licks the lick port while having a velocity of less than 0.025m/s to eliminate random licks. The mouse completes a full trial if it reaches the rewarding odor radius and licks to receive the water reward within 30s. If 30s pass before the completion of the trial then it is considered missed and a new trial starts without time delay. If the licks happen outside of the rewarding radius, then a new trial will start after a time penalty (∼1-2s). These conditions assist in task engagement, as the animal has a limited amount of time to complete a full trial and receive reward. During training, we implemented the staircase paradigm (Fig. 3c) with the increase in difficulty including reduced reward radius, increased odor diffusion throughout the virtual space and alternative initial position (e.g., start from an unrewarded odor’s radius).

### Quantification and statistical analysis

Analysis was performed using Python ≥3.8 and ≤3.12 with NumPy (Harris et al., 2020) and SciPy (Virtanen et al., 2020). Visualizations were made with Matplotlib (Hunter, 2007) and Seaborn (Waskom, 2021), and shadings represent standard error of the means.

## Supplementary Information

Supplementary Figure 1 - Synchronization metrics

Supplementary Figure 2 - Relational database schema for experiment parameters

Supplementary Figure 3 - Relational database schema for animal activity and behavior parameters

Supplementary Figure 4 - Stimulus schema for stimulus parameters

Supplementary Figure 5 - Control Table Monitor Supplementary

Figure 6 - Real time Animal Activity Monitor

Supplementary Figure 7 - Interface schema for hardware configuration Supplementary

Figure 8 - Lab Experiment Task Management

Supplementary Table 1 - Indicative component costs for one EthoPy homecage behavioral training setup with screen

Supplementary Movie 1 - Example behavioral sessions with EthoPy across systems

**Supplementary Fig. 1.**
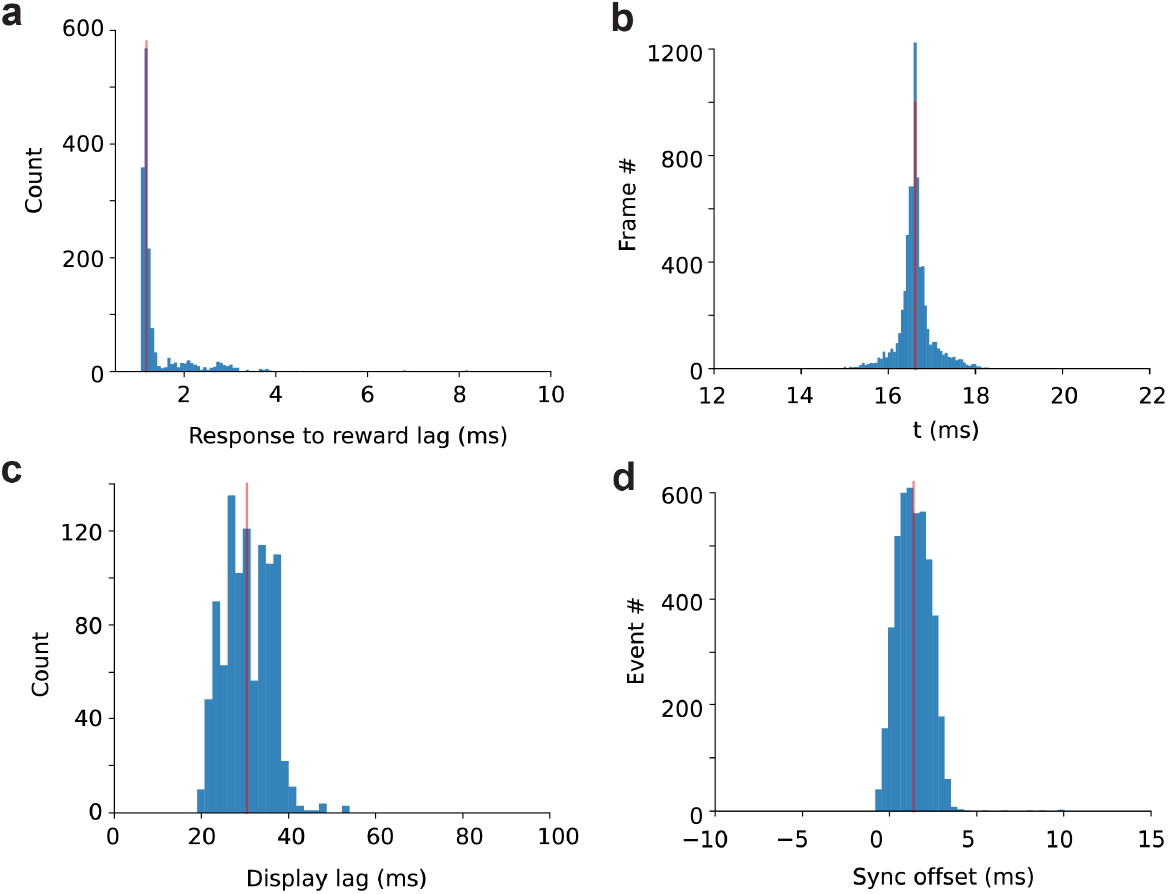
Synchronization metrics. **a**. The temporal delay between a lick response and subsequent reward delivery averaged 1.2ms, reflecting the RP’s response latency. **b**. The average delay between two subsequent frame flips of the RP 7” monitor while real-time rendering and presenting moving objects with Panda3D, was 16.6 ms. **c**. The average RP 7” monitor response lag, measured as the delay between the frame flip command and actual visual change, was 30.5ms. **d**. Using Precision Time Protocol (PTP) software-based synchronization, we estimated the average temporal offset between corresponding events logged on two separate computers to be 1.4ms, indicating a high degree of temporal alignment across systems. For details, see section Synchronization in methods.

**Supplementary Fig. 2.**
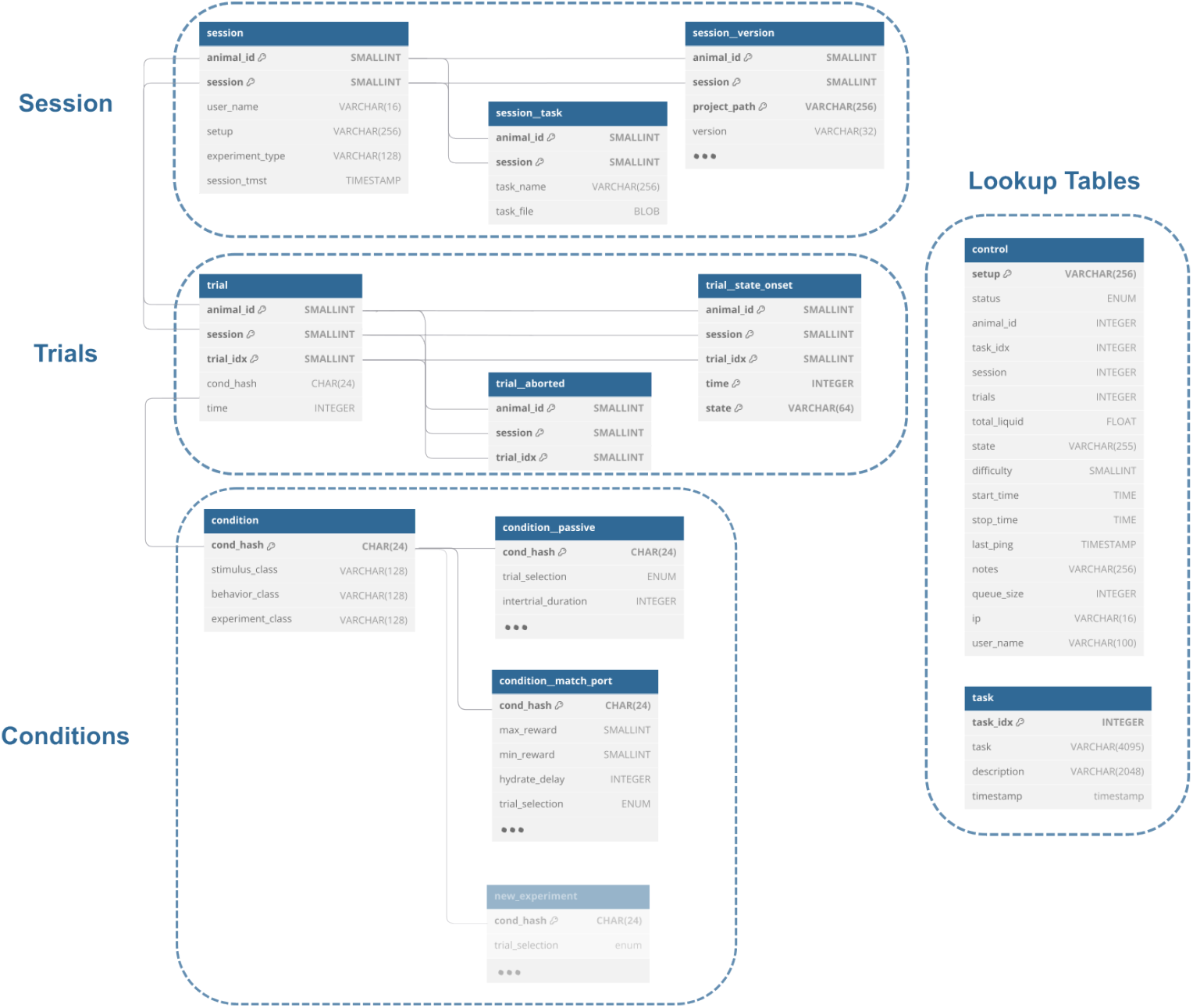
Relational database schema for experiment parameters. Each session represents a unique experimental run for an individual animal. Session metadata, including task configuration files and code versioning, is stored in the session, session_task, and session_version tables. A session is linked to its trials through shared primary keys, as shown by the connections on the left. A session’s aborted trials are logged in trial_aborted. A trial further includes the timings of its state transitions through the trial_state_onset table. Each trial is associated with the specific experiment, stimulus, and behavior classes used, via the unique cond_hash. Conditions are connected with further specialized tables (e.g., condition_passive, condition_match_port, etc.) that store experiment-specific parameters. For online monitoring, the control table tracks real-time execution status and system state, while the task table defines the available task configurations. Created using dbdiagram.io and modified by authors.

**Supplementary Fig. 3.**
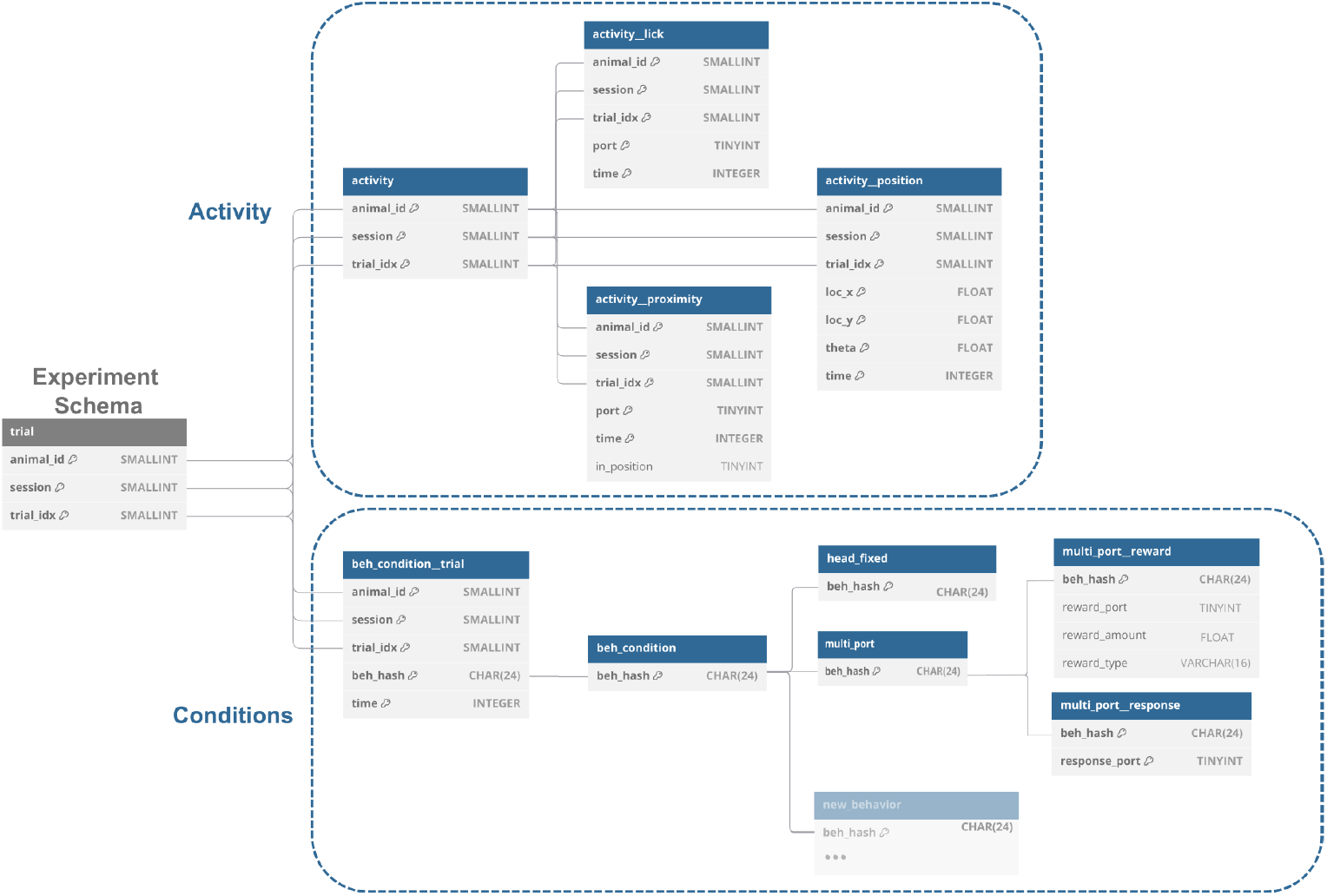
Relational database schema for animal activity and behavior parameters. All behavior-related tables are connected through shared trial keys (animal_id, session, trial_idx) with the trial table of the experiment schema. The Activity table records animal behavior per trial, including the licks, port proximity, and position. The Condition table defines trial-specific behavioral parameters linked via beh_hash, including, for example, the targeted response port and the reward delivery port. Created using dbdiagram.io and modified by authors.

**Supplementary Fig. 4.**
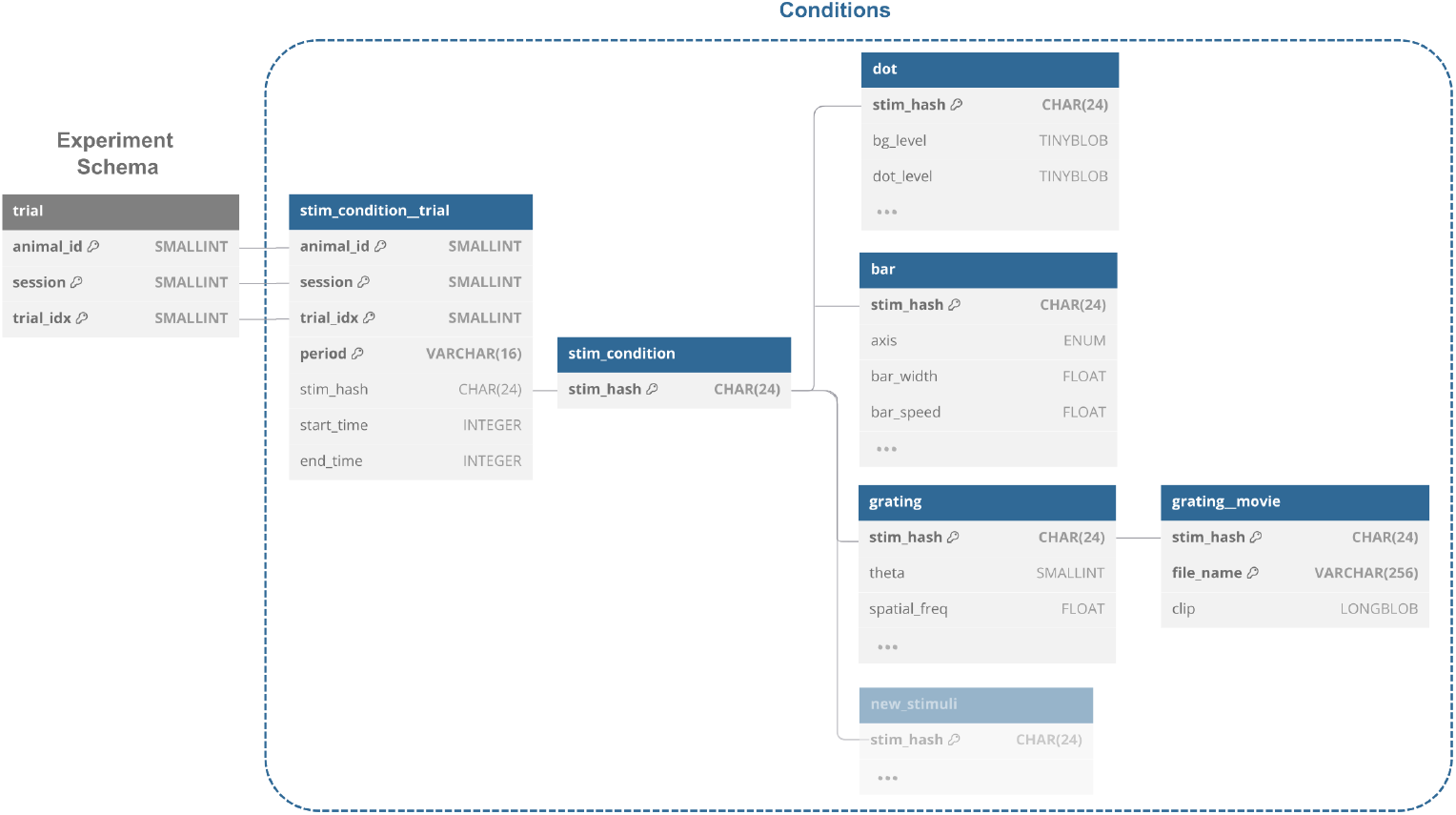
Stimulus schema for stimulus parameters. Each trial is linked to a specific stimulus configuration. The stim_condition_trial connects the stimuli to the individual trials and records presentation timings. The stim_condition table defines the stim_hash that links to the different stimuli (e.g., dot, bar, grating, grating_movie) parameters. Created using dbdiagram.io and modified by authors.

**Supplementary Fig. 5.**
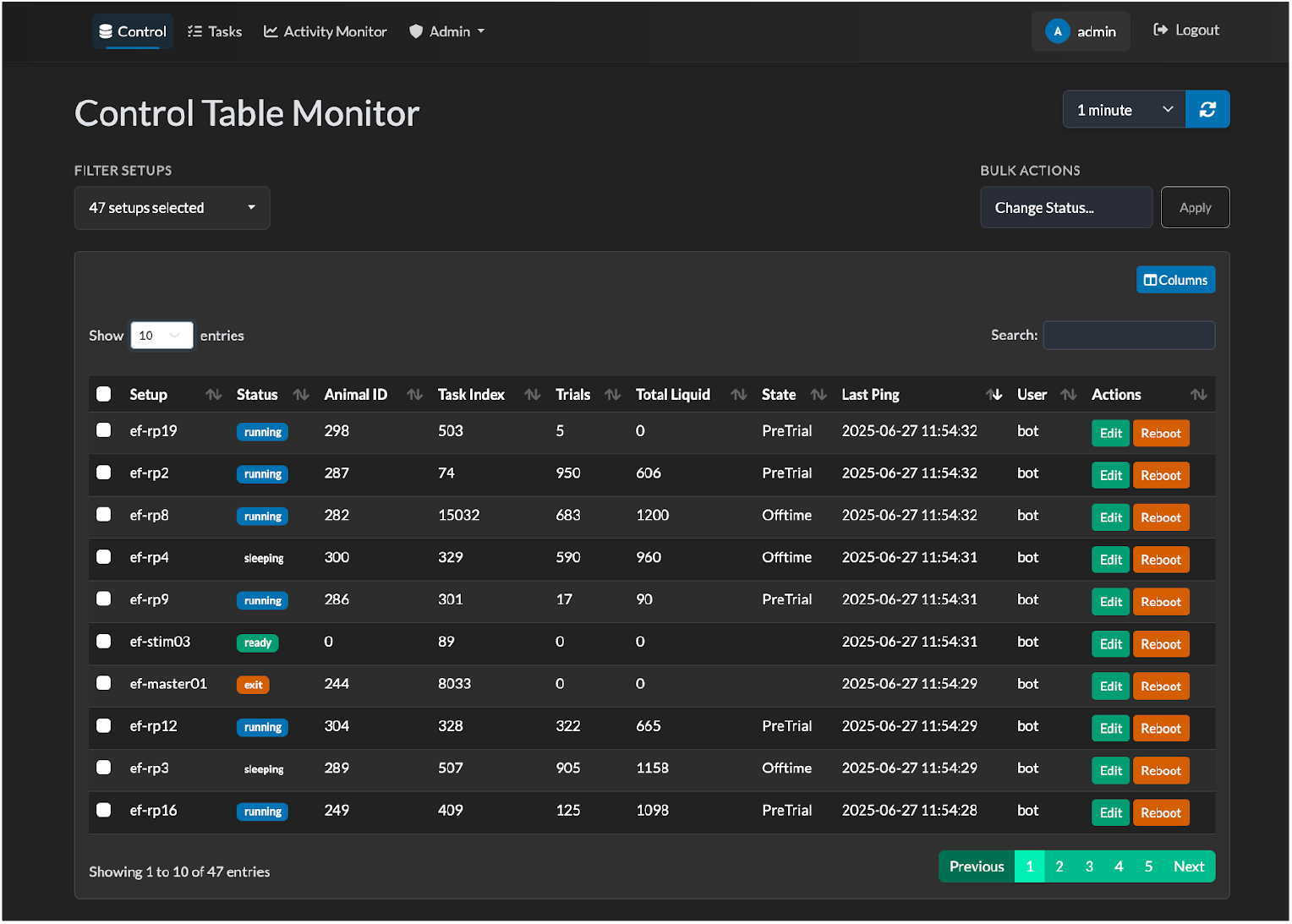
Control Table Monitor. This control panel provides an overview of the status of all experimental setups in real time. For each entry, the default columns are the setup name, operational status (e.g., running or stopped), associated Animal ID, the task index currently assigned, the last system ping (with warnings for delayed responses), and the user name. Optionally, additional parameters can be monitored, such as the current number of trials, the total liquid, and the current state of the experiment. Options to edit entry columns or reboot setups are also available. This table is essential for monitoring and controlling the experiments through the web.

**Supplementary Fig. 6.**
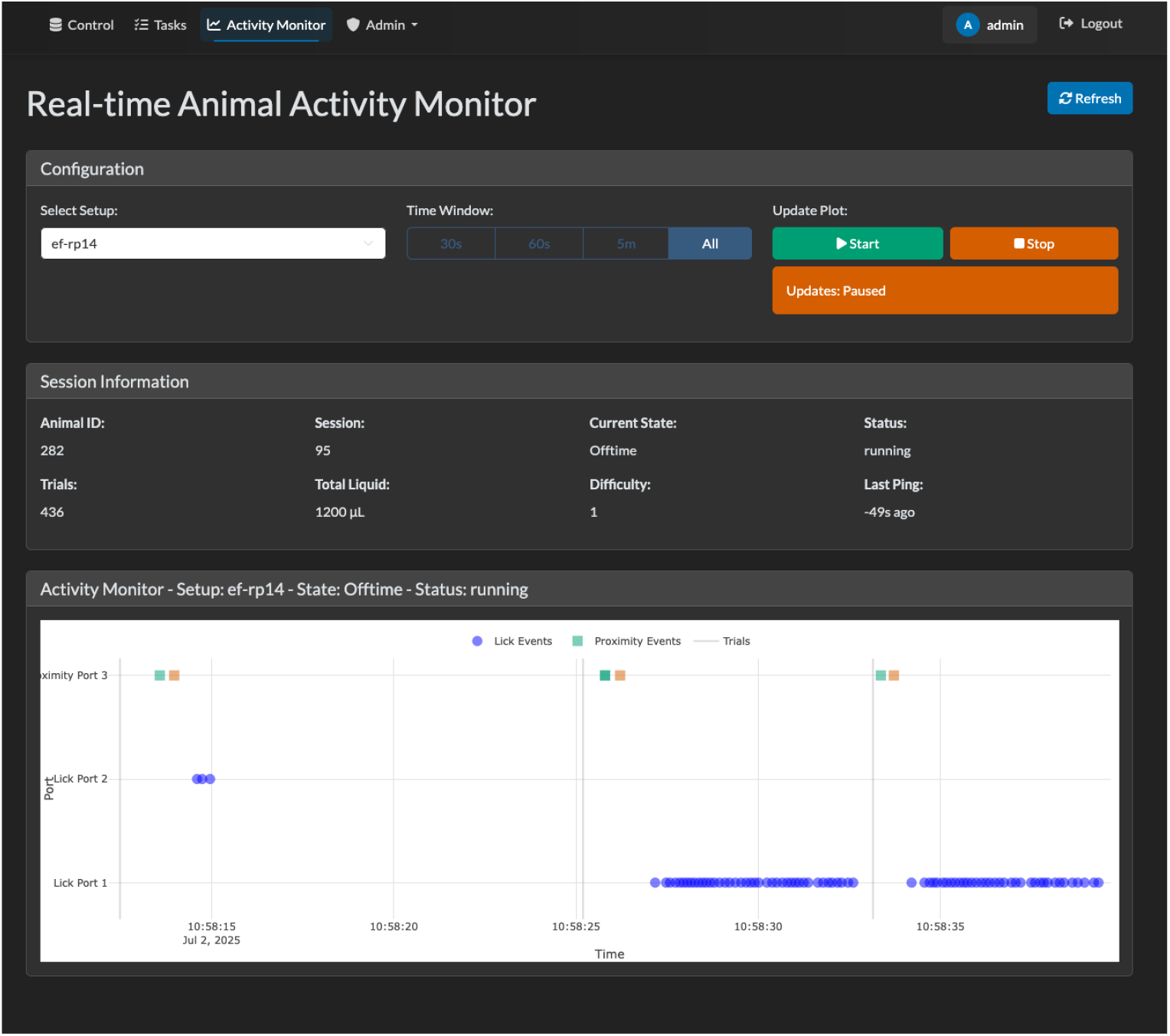
Real time Animal Activity Monitor. This panel displays the real-time monitoring used to track and visualize animals’ behavior. In the configuration panel, the user selects the setup, the displayed time window, and controls the start and stop of data streaming. Session metadata, such as Animal ID, current trial state, number of trials, etc., are also shown. A live plot illustrates behavioral events (e.g., lick and proximity events) across different ports over time and the timings of each trial start, facilitating rapid assessment of the ongoing experiment.

**Supplementary Fig. 7.**
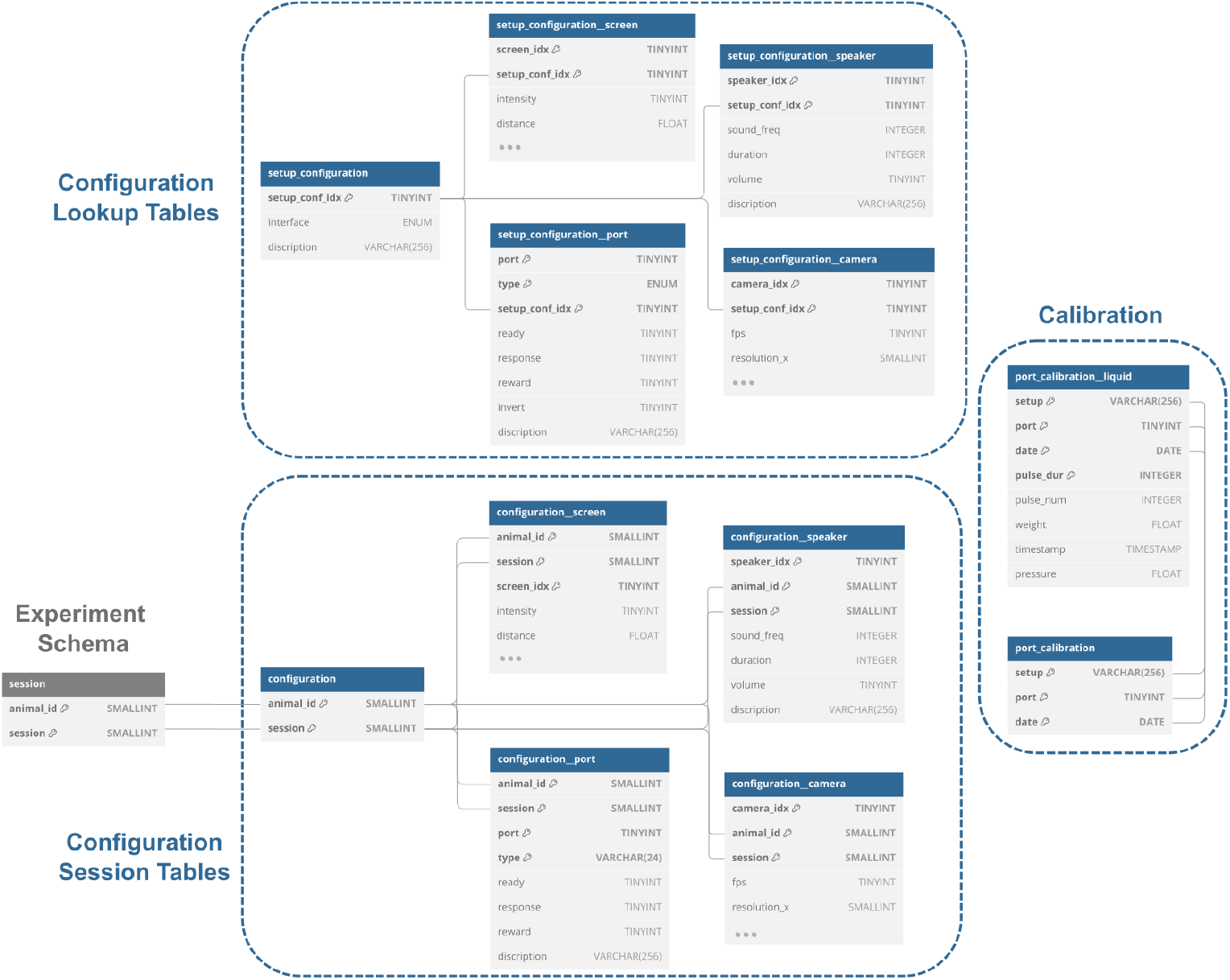
Interface schema for hardware configuration. The setup configuration tables define the parameter settings for the hardware components (e.g., ports, screens, speakers), referenced via setup_conf_idx. Session-specific configurations are stored in dedicated configuration session tables to preserve the exact setup used in each experiment. The calibration tables log port-specific measurements (e.g., liquid dispensation) to guarantee reproducibility and precision. Created using dbdiagram.io and modified by authors.

**Supplementary Fig. 8.**
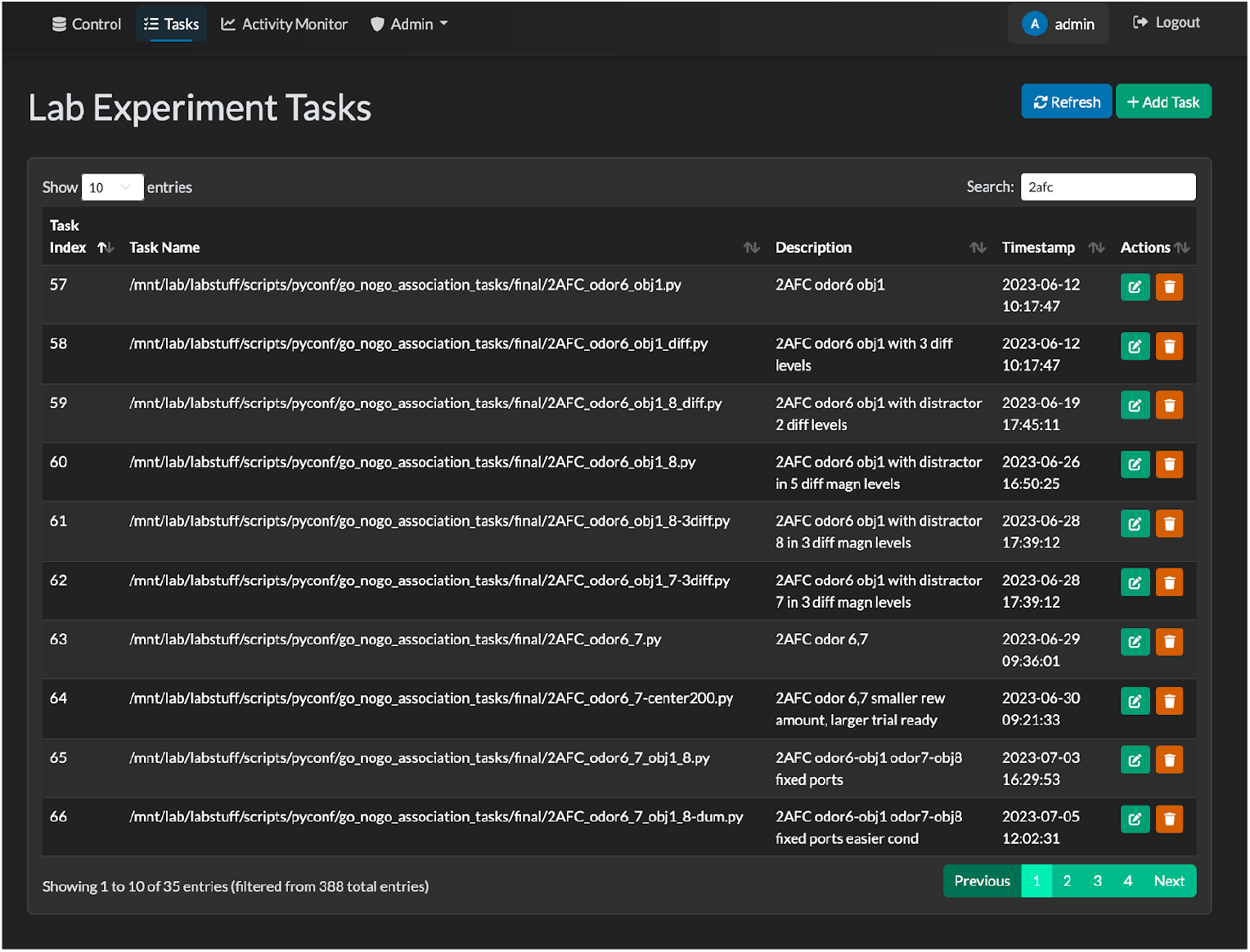
Lab Experiment Task Management. This page lists the available experimental task configuration files that can be assigned to each setup. Each row corresponds to a python task script with its index, name, description, and timestamp of creation. Users can edit or delete tasks using the corresponding action buttons. This panel enables experimenters to manage and organize the tasks.

**Supplementary Table 1.**
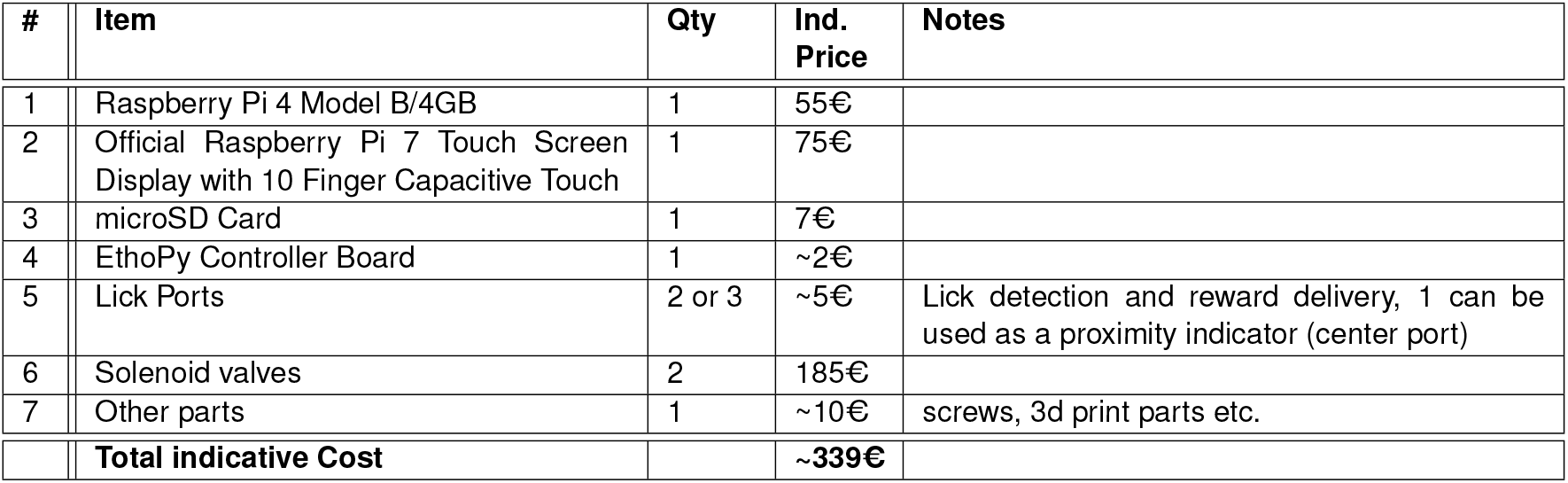
Indicative component costs for an EthoPy homecage behavioral training setup with screen. The table presents itemized costs for establishing an automated behavioral training setup capable of operating within an animal’s homecage environment. Core components include a Raspberry Pi 4 computer with screen, custom-designed controller board, and 2 or 3 lick detection ports and precision solenoid valves for reward delivery.

| # | Item | Qty | Ind. Price | Notes |
| --- | --- | --- | --- | --- |
| 1 | Raspberry Pi 4 Model B/4GB | 1 | 55€ |  |
| 2 | Official Raspberry Pi 7 Touch Screen Display with 10 Finger Capacitive Touch | 1 | 75€ |  |
| 3 | microSD Card | 1 | 7€ |  |
| 4 | EthoPy Controller Board | 1 | ~2€ |  |
| 5 | Lick Ports | 2 or 3 | ~5€ | Lick detection and reward delivery, 1 can be used as a proximity indicator (center port) |
| 6 | Solenoid valves | 2 | 185€ |  |
| 7 | Other parts | 1 | ~10€ | screws, 3d print parts etc. |
|  | **Total indicative Cost** |  | **~339€** |  |

**Supplementary Movie 1. Example behavioral sessions with EthoPy across systems** Indicative sessions across Homecage, Air-track, Spherical Treadmill and Open-field systems.

## Notes

https://github.com/ef-lab/ethopy_package

### Competing Interest Statement

The authors have declared no competing interest.

## Reference

J. J. Jun, N. A. Steinmetz, J. H. Siegle, D. J. Denman, M. Bauza, B. Barbarits, A. K. Lee, C. A. Anastassiou, A. Andrei, C. Aydın, M. Barbic, T. J. Blanche, V. Bonin, J. Couto, B. Dutta, S. L. Gratiy, D. A. Gutnisky, M. Häusser, B. Karsh, P. Ledochowitsch, C. M. Lopez, C. Mitelut, S. Musa, M. Okun, M. Pachitariu, J. Putzeys, P. D. Rich, C. Rossant, W.-l. Sun, K. Svoboda, M. Carandini, K. D. Harris, C. Koch, J. O’Keefe, and T. D. Harris. Fully integrated silicon probes for high-density recording of neural activity. Nature, 551(7679):232–236, November 2017. ISSN 0028-0836, 1476-4687. doi: 10.1038/nature24636.

N. A. Steinmetz, C. Aydin, A. Lebedeva, M. Okun, M. Pachitariu, M. Bauza, M. Beau, J. Bhagat, C. Böhm, M. Broux, S. Chen, J. Colonell, R. J. Gardner, B. Karsh, F. Kloosterman, D. Kostadinov, C. Mora-Lopez, J. O’Callaghan, J. Park, J. Putzeys, B. Sauerbrei, R. J. J. Van Daal, A. Z. Vollan, S. Wang, M. Welkenhuysen, Z. Ye, J. T. Dudman, B. Dutta, A. W. Hantman, K. D. Harris, A. K. Lee, E. I. Moser, J. O’Keefe, A. Renart, K. Svoboda, M. Häusser, S. Haesler, M. Carandini, and T. D. Harris. Neuropixels 2.0: A miniaturized high-density probe for stable, long-term brain recordings. Science, 372(6539):eabf4588, April 2021. ISSN 0036-8075, 1095-9203. doi: 10.1126/science.abf4588.

A. Lakunina, K. Z. Socha, A. Ladd, A. J. Bowen, S. Chen, J. Colonell, A. Doshi, B. Karsh, M. Krumin, P. Kulik, A. Li, P. Neutens, J. O’Callaghan, M. Olsen, J. Putzeys, H. A. Tilmans, Z. Ye, M. Welkenhuysen, M. Häusser, C. Koch, J. T. Ting, Neuropixels Opto Consortium, B. Dutta, T. D. Harris, N. A. Steinmetz, K. Svoboda, J. H. Siegle, and M. Carandini. Neuropixels Opto: Combining high-resolution electrophysiology and optogenetics, February 2025.

N. J. Sofroniew, D. Flickinger, J. King, and K. Svoboda. A large field of view two-photon mesoscope with subcellular resolution for in vivo imaging. eLife, 5:e14472, June 2016. ISSN 2050-084X. doi: 10.7554/eLife.14472.

Y. Zhang, L. Yuan, Q. Zhu, J. Wu, T. Nöbauer, R. Zhang, G. Xiao, M. Wang, H. Xie, Z. Guo, Q. Dai, and A. Vaziri. A miniaturized mesoscope for the large-scale single-neuron-resolved imaging of neuronal activity in freely behaving mice. Nature Biomedical Engineering, 8(6):754–774, June 2024. ISSN 2157-846X. doi: 10.1038/s41551-024-01226-2.

C. Guo, G. J. Blair, M. Sehgal, F. N. Sangiuliano Jimka, A. Bellafard, A. J. Silva, P. Golshani, M. A. Basso, H. T. Blair, and D. Aharoni. Miniscope-LFOV: A large-field-of-view, single-cell-resolution, miniature microscope for wired and wire-free imaging of neural dynamics in freely behaving animals. Science Advances, 9(16):eadg3918, April 2023. ISSN 2375-2548. doi: 10.1126/sciadv.adg3918.

J. W. Krakauer, A. A. Ghazanfar, A. Gomez-Marin, M. A. MacIver, and D. Poeppel. Neuroscience Needs Behavior: Correcting a Reductionist Bias. Neuron, 93(3):480–490, February 2017. ISSN 08966273. doi: 10.1016/j.neuron.2016.12.041.

S. M. Peters, H. H. Pothuizen, and B. M. Spruijt. Ethological concepts enhance the translational value of animal models. European Journal of Pharmacology, 759:42–50, July 2015. ISSN 00142999. doi: 10.1016/j.ejphar.2015.03.043.

C. Vielle. Beyond the Illusion of Controlled Environments: How to Embrace Ecological Pertinence in Research? European Journal of Neuroscience, 61(1):e16661, January 2025. ISSN 0953-816X, 1460-9568. doi: 10.1111/ejn.16661.

Y. Shemesh and A. Chen. A paradigm shift in translational psychiatry through rodent neuroethology. Molecular Psychiatry, 28(3):993–1003, March 2023. ISSN 1359-4184, 1476-5578. doi: 10.1038/s41380-022-01913-z.

A. Nasr, S. E. Dominiak, K. Sehara, M. A. Nashaat, R. N. S. Sachdev, and M. E. Larkum. Efficient training approaches for optimizing behavioral performance and reducing head fixation time. PLOS ONE, 17(11):e0276531, November 2022. ISSN 1932-6203. doi: 10.1371/journal.pone.0276531.

M. J. Castelhano-Carlos, V. Baumans, and N. Sousa. PhenoWorld: addressing animal welfare in a new paradigm to house and assess rat behaviour. Laboratory Animals, 51(1):36–43, February 2017. ISSN 0023-6772, 1758-1117. doi: 10.1177/0023677216638642. Publisher: SAGE Publications.

M. R. Watson, B. Voloh, C. Thomas, A. Hasan, and T. Womelsdorf. USE: An integrative suite for temporally-precise psychophysical experiments in virtual environments for human, nonhuman, and artificially intelligent agents. Journal of Neuroscience Methods, 326:108374, October 2019. ISSN 0165-0270. doi: 10.1016/j.jneumeth.2019.108374. Publisher: Elsevier BV.

Nourizonoz, R. Zimmermann, C. L. A. Ho, S. Pellat, Y. Ormen, C. Prévost-Solié, G. Reymond, F. Pifferi, F. Aujard, A. Herrel, and D. Huber. EthoLoop: automated closed-loop neuroethology in naturalistic environments. Nature Methods, 17(10):1052–1059, October 2020. ISSN 1548-7091, 1548-7105. doi: 10.1038/s41592-020-0961-2. Publisher: Springer Science and Business Media LLC.

Kim, S. C. Kenchappa, A. Sunkara, T.-Y. Chang, L. Thompson, R. Doudlah, and A. Rosenberg. Real-time experimental control using network-based parallel processing. eLife, 8, February 2019. ISSN 2050-084X. doi: 10.7554/elife.40231. Publisher: eLife Sciences Publications, Ltd.

G. Lopes, N. Bonacchi, J. FrazÃ£o, J. P. Neto, B. V. Atallah, S. Soares, L. Moreira, S. Matias, P. M. Itskov, P. A. Correia, R. E. Medina, L. Calcaterra, E. Dreosti, J. J. Paton, and A. R. Kampff. Bonsai: an event-based framework for processing and controlling data streams. Frontiers in Neuroinformatics, 9, April 2015. ISSN 1662-5196. doi: 10.3389/fninf.2015.00007.

S. J. Sukoff Rizzo and J. L. Silverman. Methodological Considerations for Optimizing and Validating Behavioral Assays. Current Protocols in Mouse Biology, 6(4):364–379, December 2016. ISSN 2161-2617, 2161-2617. doi: 10.1002/cpmo.17.

D. Zoccolan and A. Di Filippo. Methodological Approaches to the Behavioural Investigation of Visual Perception in Rodents. In Handbook of Behavioral Neuroscience, volume 27, pages 69–101. Elsevier, 2018. ISBN 978-0-12-812012-5. doi: 10.1016/B978-0-12-812012-5.00005-7.

E. J. Chesler, S. G. Wilson, W. R. Lariviere, S. L. Rodriguez-Zas, and J. S. Mogil. Identification and ranking of genetic and laboratory environment factors influencing a behavioral trait, thermal nociception, via computational analysis of a large data archive. Neuroscience & Biobehavioral Reviews, 26(8):907–923, December 2002. ISSN 01497634. doi: 10.1016/S0149-7634(02)00103-3.

R. E. Sorge, L. J. Martin, K. A. Isbester, S. G. Sotocinal, S. Rosen, A. H. Tuttle, J. S. Wieskopf, E. L. Acland, A. Dokova, B. Kadoura, P. Leger, J. C. S. Mapplebeck, M. McPhail, A. Delaney, G. Wigerblad, A. P. Schumann, T. Quinn, J. Frasnelli, C. I. Svensson, W. F. Sternberg, and J. S. Mogil. Olfactory exposure to males, including men, causes stress and related analgesia in rodents. Nature Methods, 11(6):629–632, June 2014. ISSN 1548-7091, 1548-7105. doi: 10.1038/nmeth.2935.

J. P. Balcombe, N. D. Barnard, and C. Sandusky. Laboratory routines cause animal stress. Contemporary Topics in Laboratory Animal Science, 43(6):42–51, November 2004. ISSN 1060-0558.

S. H. Richter. Automated Home-Cage Testing as a Tool to Improve Reproducibility of Behavioral Research? Frontiers in Neuroscience, 14:383, April 2020. ISSN 1662-453X. doi: 10.3389/fnins.2020.00383.

M. Nigri, J. Åhlgren, D. P. Wolfer, and V. Voikar. Role of Environment and Experimenter in Reproducibility of Behavioral Studies With Laboratory Mice. Frontiers in Behavioral Neuroscience, 16:835444, February 2022. ISSN 1662-5153. doi: 10.3389/fnbeh.2022.835444.

M. Gulinello, H. A. Mitchell, Q. Chang, W. Timothy O’Brien, Z. Zhou, T. Abel, L. Wang, J. G. Corbin, S. Veeraragavan, R. C. Samaco, N. A. Andrews, M. Fagiolini, T. B. Cole, T. M. Burbacher, and J. N. Crawley. Rigor and reproducibility in rodent behavioral research. Neurobiology of Learning and Memory, 165:106780, November 2019. ISSN 10747427. doi: 10.1016/j.nlm.2018.01.001.

A. Berditchevskaia, R. D. Cazé, and S. R. Schultz. Performance in a GO/NOGO perceptual task reflects a balance between impulsive and instrumental components of behaviour. Scientific Reports, 6(1):27389, June 2016. ISSN 2045-2322. doi: 10.1038/srep27389.

D. Hulsey, K. Zumwalt, L. Mazzucato, D. A. McCormick, and S. Jaramillo. Decision-making dynamics are predicted by arousal and uninstructed movements. Cell Reports, 43(2):113709, February 2024. ISSN 22111247. doi: 10.1016/j.celrep.2024.113709.

S. Pisupati, L. Chartarifsky-Lynn, A. Khanal, and A. K. Churchland. Lapses in perceptual decisions reflect exploration. eLife, 10:e55490, January 2021. ISSN 2050-084X. doi: 10.7554/eLife.55490.

J. F. Schweihoff, M. Loshakov, I. Pavlova, L. Kück, L. A. Ewell, and M. K. Schwarz. DeepLab-Stream enables closed-loop behavioral experiments using deep learning-based markerless, real-time posture detection. Communications Biology, 4(1):130, January 2021. ISSN 2399-3642. doi: 10.1038/s42003-021-01654-9.

T. Akam, A. Lustig, J. M. Rowland, S. K. Kapanaiah, J. Esteve-Agraz, M. Panniello, C. Márquez, M. M. Kohl, D. Kätzel, R. M. Costa, and M. E. Walton. Open-source, Python-based, hardware and software for controlling behavioural neuroscience experiments. eLife, 11:e67846, January 2022. ISSN 2050-084X. doi: 10.7554/eLife.67846.

J. L. Saunders, L. A. Ott, and M. Wehr. AUTOPILOT: Automating experiments with lots of Raspberry Pis, October 2019.

M. E. Martone. The past, present and future of neuroscience data sharing: a perspective on the state of practices and infrastructure for FAIR. Frontiers in Neuroinformatics, 17:1276407, January 2024. ISSN 1662-5196. doi: 10.3389/fninf.2023.1276407.

K. Ameen-Ali, M. Eacott, and A. Easton. A new behavioural apparatus to reduce animal numbers in multiple types of spontaneous object recognition paradigms in rats. Journal of Neuroscience Methods, 211(1):66–76, October 2012. ISSN 01650270. doi: 10.1016/j.jneumeth.2012.08.006.

R. Poddar, R. Kawai, and B. P. Ölveczky. A Fully Automated High-Throughput Training System for Rodents. PLoS ONE, 8(12):e83171, December 2013. ISSN 1932-6203. doi: 10.1371/journal.pone.0083171.

R. Aoki, T. Tsubota, Y. Goya, and A. Benucci. An automated platform for high-throughput mouse behavior and physiology with voluntary head-fixation. Nature Communications, 8(1):1196, October 2017. ISSN 2041-1723. doi: 10.1038/s41467-017-01371-0.

A. Gordon-Fennell, J. M. Barbakh, M. T. Utley, S. Singh, P. Bazzino, R. Gowrishankar, M. R. Bruchas, M. F. Roitman, and G. D. Stuber. An open-source platform for head-fixed operant and consummatory behavior. eLife, 12:e86183, August 2023. ISSN 2050-084X. doi: 10.7554/eLife.86183.

I. N. Iman, N. A. M. Yusof, U. N. Talib, N. A. Z. Ahmad, A. Norazit, J. Kumar, M. Z. Mehat, N. Jayabalan, S. Muthuraju, M. Stefaniuk, L. Kaczmarek, and M. Muzaimi. The IntelliCage System: A Review of Its Utility as a Novel Behavioral Platform for a Rodent Model of Substance Use Disorder. Frontiers in Behavioral Neuroscience, 15:683780, June 2021. ISSN 1662-5153. doi: 10.3389/fnbeh.2021.683780.

D. Yatsenko, E. Y. Walker, and A. S. Tolias. DataJoint: A Simpler Relational Data Model, 2018. Version Number: 1.

E. C. Johnson, T. T. Nguyen, B. K. Dichter, F. Zappulla, M. Kosma, K. Gunalan, Y. O. Halchenko, S. Q. Neufeld, K. Ratan, N. J. Edwards, S. Ressl, S. R. Heilbronner, M. Schirner, P. Ritter, B. Wester, S. Ghosh, M. E. Martone, F. Pestilli, and D. Yatsenko. SciOps: Achieving Productivity and Reliability in Data-Intensive Research, November 2024. 2401.00077 [q-bio].

E. Muller, J. A. Bednar, M. Diesmann, M.-O. Gewaltig, M. Hines, and A. P. Davison. Python in neuroscience. Frontiers in Neuroinformatics, 9, April 2015. ISSN 1662-5196. doi: 10.3389/fninf.2015.00011.

M. Goslin and M. Mine. The Panda3D graphics engine. Computer, 37(10):112–114, October 2004. ISSN 1558-0814. doi: 10.1109/MC.2004.180. Conference Name:Computer.

A. Mathis, P. Mamidanna, K. M. Cury, T. Abe, V. N. Murthy, M. W. Mathis, and M. Bethge. DeepLabCut: markerless pose estimation of user-defined body parts with deep learning. Nature Neuroscience, 21(9):1281–1289, September 2018. ISSN 1097-6256, 1546-1726. doi: 10.1038/s41593-018-0209-y.

G. A. Kane, G. Lopes, J. L. Saunders, A. Mathis, and M. W. Mathis. Real-time, low-latency closed-loop feedback using markerless posture tracking. eLife, 9:e61909, December 2020. ISSN 2050-084X. doi: 10.7554/eLife.61909.

J. W. Peirce. PsychoPy—Psychophysics software in Python. Journal of Neuroscience Methods, 162(1-2):8–13, May 2007. ISSN 01650270. doi: 10.1016/j.jneumeth.2006.11.017.

J. W. Peirce. Generating stimuli for neuroscience using PsychoPy. Frontiers in Neuroinformatics, 2, 2008. ISSN 16625196. doi: 10.3389/neuro.11.010.2008.

J. H. Siegle, A. C. López, Y. A. Patel, K. Abramov, S. Ohayon, and J. Voigts. Open Ephys: an open-source, plugin-based platform for multichannel electrophysiology. Journal of Neural Engineering, 14(4):045003, August 2017. ISSN 1741-2560, 1741-2552. doi: 10.1088/1741-2552/aa5eea.

E. Gamma, R. Helm, R. Johnson, and J. Vlissides. Design Patterns: Elements of Reusable Object-Oriented Software. Addison-Wesley Professional Computing Series. Pearson Education, 1994. ISBN 978-0-321-70069-8.

A. S. Tanenbaum and H. Bos. Modern operating systems. Prentice Hall, Boston, 4. ed edition, 2015. ISBN 978-0-13-359162-0 978-1-292-06142-9.

J. Teeters, K. Godfrey, R. Young, C. Dang, C. Friedsam, B. Wark, H. Asari, S. Peron, N. Li, A. Peyrache, G. Denisov, J. Siegle, S. Olsen, C. Martin, M. Chun, S. Tripathy, T. Blanche, K. Harris, G. Buzsáki, C. Koch, M. Meister, K. Svoboda, and F. Sommer. Neurodata Without Borders: Creating a Common Data Format for Neurophysiology. Neuron, 88(4):629–634, November 2015. ISSN 08966273. doi: 10.1016/j.neuron.2015.10.025.

O. Rübel, A. Tritt, R. Ly, B. K. Dichter, S. Ghosh, L. Niu, P. Baker, I. Soltesz, L. Ng, K. Svoboda, L. Frank, and K. E. Bouchard. The Neurodata Without Borders ecosystem for neurophysiological data science. eLife, 11:e78362, October 2022. ISSN 2050-084X. doi: 10.7554/eLife.78362.

M. A. Nashaat, H. Oraby, R. N. S. Sachdev, Y. Winter, and M. E. Larkum. Air-Track: a real-world floating environment for active sensing in head-fixed mice. Journal of Neurophysiology, 116 (4):1542–1553, October 2016. ISSN 0022-3077, 1522-1598. doi: 10.1152/jn.00088.2016.

N. M. Abraham, D. Guerin, K. Bhaukaurally, and A. Carleton. Similar Odor Discrimination Behavior in Head-Restrained and Freely Moving Mice. PLoS ONE, 7(12):e51789, December 2012. ISSN 1932-6203. doi: 10.1371/journal.pone.0051789.

S. Ceballo, J. Bourg, A. Kempf, Z. Piwkowska, A. Daret, P. Pinson, T. Deneux, S. Rumpel, and B. Bathellier. Cortical recruitment determines learning dynamics and strategy. Nature Communications, 10(1):1479, April 2019. ISSN 2041-1723. doi: 10.1038/s41467-019-09450-0.

P. A. Dudchenko. An overview of the tasks used to test working memory in rodents. Neuroscience & Biobehavioral Reviews, 28(7):699–709, January 2004. ISSN 0149-7634. doi: 10.1016/j.neubiorev.2004.09.002. Publisher: Elsevier BV.

J. R. Gibson and J. H. R. Maunsell. Sensory Modality Specificity of Neural Activity Related to Memory in Visual Cortex. Journal of Neurophysiology, 78(3):1263–1275, September 1997. ISSN 0022-3077, 1522-1598. doi: 10.1152/jn.1997.78.3.1263. Publisher: American Physiological Society.

E. Froudarakis, U. Cohen, M. Diamantaki, E. Y. Walker, J. Reimer, P. Berens, H. Sompolinsky, and A. S. Tolias. Object manifold geometry across the mouse cortical visual hierarchy, August 2020.

K. M. Roddick, H. M. Schellinck, and R. E. Brown. Olfactory delayed matching to sample performance in mice: Sex differences in the 5XFAD mouse model of Alzheimer’s disease. Behavioural Brain Research, 270:165–170, August 2014. ISSN 0166-4328. doi: 10.1016/j.bbr.2014.04.038. Publisher: Elsevier BV.

Y. Sakurai. Hippocampal cells have behavioral correlates during the performance of an auditory working memory task in the rat. Behavioral Neuroscience, 104(2):253–263, April 1990. ISSN 0735-7044. doi: 10.1037//0735-7044.104.2.253.

J. H. Sul, S. Jo, D. Lee, and M. W. Jung. Role of rodent secondary motor cortex in value-based action selection. Nature Neuroscience, 14(9):1202–1208, September 2011. ISSN 1097-6256, 1546-1726. doi: 10.1038/nn.2881.

C. Findling, F. Hubert, International Brain Laboratory, L. Acerbi, B. Benson, J. Benson, D. Birman, N. Bonacchi, S. Bruijns, M. Carandini, J. A. Catarino, G. A. Chapuis, A. K. Churchland, Y. Dan, F. Davatolhagh, E. E. DeWitt, T. A. Engel, M. Fabbri, M. Faulkner, I. R. Fiete, L. Freitas-Silva, B. Gerçek, K. D. Harris, M. Häusser, S. B. Hofer, F. Hu, J. M. Huntenburg, A. Khanal, C. Krasniak, C. Langdon, P. E. Latham, P. Y. P. Lau, Z. Mainen, G. T. Meijer, N. J. Miska, T. D. Mrsic-Flogel, J.-P. Noel, K. Nylund, A. Pan-Vazquez, L. Paninski, J. Pillow, C. Rossant, N. Roth, R. Schaeffer, M. Schartner, Y. Shi, K. Z. Socha, N. A. Steinmetz, K. Svoboda, C. Tessereau, A. E. Urai, M. J. Wells, S. J. West, M. R. Whiteway, O. Winter, I. B. Witten, A. Zador, Y. Zhang, P. Dayan, and A. Pouget. Brain-wide representations of prior information in mouse decision-making, July 2023.

C. M. Niell and M. P. Stryker. Modulation of Visual Responses by Behavioral State in Mouse Visual Cortex. Neuron, 65(4):472–479, February 2010. ISSN 08966273. doi: 10.1016/j.neuron.2010.01.033.

A. B. Saleem, A. Ayaz, K. J. Jeffery, K. D. Harris, and M. Carandini. Integration of visual motion and locomotion in mouse visual cortex. Nature Neuroscience, 16(12):1864–1869, December 2013. ISSN 1097-6256, 1546-1726. doi: 10.1038/nn.3567.

D. A. Dombeck, C. D. Harvey, L. Tian, L. L. Looger, and D. W. Tank. Functional imaging of hippocampal place cells at cellular resolution during virtual navigation. Nature Neuroscience, 13(11):1433–1440, November 2010. ISSN 1097-6256, 1546-1726. doi: 10.1038/nn.2648.

C. Domnisoru, A. A. Kinkhabwala, and D. W. Tank. Membrane potential dynamics of grid cells. Nature, 495(7440):199–204, March 2013. ISSN 0028-0836, 1476-4687. doi: 10.1038/nature11973.

R. Gau, S. Noble, K. Heuer, K. L. Bottenhorn, I. P. Bilgin, Y.-F. Yang, J. M. Huntenburg, J. M. Bayer, R. A. Bethlehem, S. A. Rhoads, C. Vogelbacher, V. Borghesani, E. Levitis, H.-T. Wang, S. Van Den Bossche, X. Kobeleva, J. H. Legarreta, S. Guay, S. M. Atay, G. P. Varoquaux, D. C. Huijser, M. S. Sandström, P. Herholz, S. A. Nastase, A. Badhwar, G. Dumas, S. Schwab, S. Moia, M. Dayan, Y. Bassil, P. P. Brooks, M. Mancini,. M. Shine, D. O’Connor, X. Xie, D. Poggiali, P. Friedrich, A. S. Heinsfeld, L. Riedl, R. Toro, C. Caballero-Gaudes, A. Eklund, K. G. Garner, C. R. Nolan, D. V. Demeter, F. A. Barrios, J. S. Merchant, E. A. McDevitt, R. Oostenveld, R. C. Craddock, A. Rokem, A. Doyle, S. S. Ghosh, A. Nikolaidis, O. W. Stanley, E. Uruñuela, N. Anousheh, A. Arnatkeviciute, G. Auzias, D. Bachar, E. Bannier, R. Basanisi, A. Basavaraj, M. Bedini, P. Bellec, R. A. Benn, K. Berluti, S. Bollmann, S. Bollmann, C. Bradley, J. Brown, A. Buchweitz, P. Callahan, M. Y. Chan, B. Q. Chandio, T. Cheng, S. Chopra, A. W. Chung, T. G. Close, E. Combrisson, G. Cona, R. T. Constable, C. Cury, K. Dadi, P. F. Damasceno, S. Das, F. De Vico Fallani, K. DeStasio, E. W. Dickie, L. Dorfschmidt, E. P. Duff, E. DuPre, S. Dziura, N. B. Esper, O. Esteban, S. Fadnavis, G. Flandin, J. E. Flannery, J. Flournoy, S. J. Forkel, A. R. Franco, S. Ganesan, S. Gao, J. C. García Alanis, E. Garyfallidis, T. Glatard, E. Glerean, J. Gonzalez-Castillo, C. D. Gould Van Praag, A. S. Greene, G. Gupta, C. A. Hahn, Y. O. Halchenko, D. Handwerker, T. S. Hartmann, V. Hayot-Sasson, S. Heunis, F. Hoffstaedter, D. M. Hohmann, C. Horien, H.-I. Ioanas, A. Iordan, C. Jiang, M. Joseph, J. Kai, A. Karakuzu, D. N. Kennedy, A. Keshavan, A. R. Khan, G. Kiar, P. C. Klink, V. Koppelmans, S. Koudoro, A. R. Laird, G. Langs, M. Laws, R. Licandro, S.-L. Liew, T. Lipic, K. Litinas, D. J. Lurie, D. Lussier, C. R. Madan, L.-T. Mais, S. Mansour L, J. Manzano-Patron, D. Maoutsa, M. Marcon, D. S. Margulies, G. Marinato, D. Marinazzo, C. J. Markiewicz, C. Maumet, F. Meneguzzi, D. Meunier, M. P. Milham, K. L. Mills, D. Momi, C. A. Moreau, A. Motala, I. Moxon-Emre, T. E. Nichols, D. M. Nielson, G. Nilsonne, L. Novello, C. O’Brien, E. Olafson, L. D. Oliver, J. A. Onofrey, E. R. Orchard, K. Oudyk, P. J. Park, M. Parsapoor, L. Pasquini, S. Peltier, C. R. Pernet, R. Pienaar, P. Pinheiro-Chagas, J.-B. Poline, A. Qiu, T. Quendera, L. C. Rice, J. Rocha-Hidalgo, S. Rutherford, M. Scharinger, D. Scheinost, D. Shariq, T. B. Shaw, V. Siless, M. Simmonite, N. Sirmpilatze, H. Spence, J. Sprenger, A. Stajduhar, M. Szinte, S. Takerkart, A. Tam, L. Tejavibulya, M. Thiebaut De Schotten, I. Thome, L. Tomaz Da Silva, N. Traut, L. Q. Uddin, A. Vallesi, J. W. VanMeter, N. Vijayakumar, M. V. Di Oleggio Castello, J. Vohryzek, J. Vukojević, K. J. Whitaker, L. Whitmore, S. Wideman, S. T. Witt, H. Xie, T. Xu, C.-G. Yan, F.-C. Yeh, B. T. Yeo, and X.-N. Zuo. Brainhack: Developing a culture of open, inclusive, community-driven neuroscience. Neuron, 109(11):1769–1775, June 2021. ISSN 08966273. doi: 10.1016/j.neuron.2021.04.001.

M. Diamantaki and A. Papoutsi. Gather your neurons and model together: Community times ahead. PLOS Biology, 22(11):e3002839, November 2024. ISSN 1545-7885. doi: 10.1371/journal.pbio.3002839.

C. V. Vorhees and M. T. Williams. Morris water maze: procedures for assessing spatial and related forms of learning and memory. Nature Protocols, 1(2):848–858, August 2006. ISSN 1754-2189, 1750-2799. doi: 10.1038/nprot.2006.116.

M. Nashaat, H. Oraby, F. Krasniqi, S. T. Goh-Sauerbier, M. Bosc, S. Koerner, S. Karayel, Kepecs, and M. Larkum. The neural mechanisms of fast versus slow decision-making, August 2024.

reutwein. Adaptive psychophysical procedures. Vision Research, 35(17):2503–2522, September 1995. ISSN 0042-6989.

M. A. Garcıa-Perez. Forced-choice staircases with fixed step sizes: asymptotic and small-sample properties. Vision Research, 38(12):1861–1881, June 1998. ISSN 0042-6989. doi: 10.1016/s0042-6989(97)00340-4. Publisher: Elsevier BV.

M. R. Leek. Adaptive procedures in psychophysical research. Perception & Psychophysics, 63 (8):1279–1292, November 2001. ISSN 0031-5117, 1532-5962. doi: 10.3758/bf03194543. Publisher: Springer Science and Business Media LLC.

C. R. Harris, K. J. Millman, S. J. Van Der Walt, R. Gommers, P. Virtanen, D. Cournapeau, E. Wieser, J. Taylor, S. Berg, N. J. Smith, R. Kern, M. Picus, S. Hoyer, M. H. Van Kerkwijk, M. Brett, A. Haldane, J. F. Del Río, M. Wiebe, P. Peterson, P. Gérard-Marchant, K. Sheppard, T. Reddy, W. Weckesser, H. Abbasi, C. Gohlke, and T. E. Oliphant. Array programming with NumPy. Nature, 585(7825):357–362, September 2020. ISSN 0028-0836, 1476-4687. doi: 10.1038/s41586-020-2649-2. Publisher: Springer Science and Business Media LLC.

P. Virtanen, R. Gommers, T. E. Oliphant, M. Haberland, T. Reddy, D. Cournapeau, E. Burovski, P. Peterson, W. Weckesser, J. Bright, S. J. Van Der Walt, M. Brett, J. Wilson, K. J. Millman, N. Mayorov, A. R. J. Nelson, E. Jones, R. Kern, E. Larson, C. J. Carey, I. Polat, Y. Feng, E. W. Moore, J. VanderPlas, D. Laxalde, J. Perktold, R. Cimrman, I. Henriksen, E. A. Quintero, C. R. Harris, A. M. Archibald, A. H. Ribeiro, F. Pedregosa, P. Van Mulbregt, SciPy 1.0 Contributors, A. Vijaykumar, A. P. Bardelli, A. Rothberg, A. Hilboll, A. Kloeckner, A. Scopatz, A. Lee, A. Rokem, C. N. Woods, C. Fulton, C. Masson, C. Häggström, C. Fitzgerald, D. A. Nicholson, D. R. Hagen, D. V. Pasechnik, E. Olivetti, E. Martin, E. Wieser, F. Silva, F. Lenders, F. Wilhelm, G. Young, G. A. Price, G.-L. Ingold, G. E. Allen, G. R. Lee, H. Audren, I. Probst, J. P. Dietrich, J. Silterra, J. T. Webber, J. Slavič, J. Nothman, J. Buchner, J. Kulick, J. L. Schönberger, J. V. De Miranda Cardoso, J. Reimer, J. Harrington, J. L. C. Rodríguez, J. Nunez-Iglesias, J. Kuczynski, K. Tritz, M. Thoma, M. Newville, M. Kümmerer, M. Bolingbroke, M. Tartre, M. Pak, N. J. Smith, N. Nowaczyk, N. Shebanov, O. Pavlyk, P. A. Brodtkorb, P. Lee, R. T. McGibbon, R. Feldbauer, S. Lewis, S. Tygier, S. Sievert, S. Vigna, S. Peterson, S. More, T. Pudlik, T. Oshima, T. J. Pingel, T. P. Robitaille, T. Spura, T. R. Jones, T. Cera, T. Leslie, T. Zito, T. Krauss, U. Upadhyay, Y. O. Halchenko, and Y. Vázquez-Baeza. SciPy 1.0: fundamental algorithms for scientific computing in Python. Nature Methods, 17(3):261–272, March 2020. ISSN 1548-7091, 1548-7105. doi: 10.1038/s41592-019-0686-2. Publisher: Springer Science and Business Media LLC.

J. D. Hunter. Matplotlib: A 2D Graphics Environment. Computing in Science & Engineering, 9(3): 90–95, 2007. ISSN 1521-9615. doi: 10.1109/mcse.2007.55. Publisher: Institute of Electrical and Electronics Engineers (IEEE).

M. Waskom. seaborn: statistical data visualization. Journal of Open Source Software, 6(60): 3021, April 2021. ISSN 2475-9066. doi: 10.21105/joss.03021. Publisher: The Open Journal.

